# Free and interfacial boundaries in individual-based models of multicellular biological systems

**DOI:** 10.1101/2022.12.13.520331

**Authors:** Domenic P. J. Germano, Adriana Zanca, Stuart T. Johnston, Jennifer A. Flegg, James M. Osborne

## Abstract

Coordination of cell behaviour is key to a myriad of biological processes including tissue morphogenesis, wound healing, and tumour growth. As such, individual-based computational models, which explicitly describe inter-cellular interactions, are commonly used to model collective cell dynamics. However, when using individual-based models, it is unclear how descriptions of cell boundaries affect overall population dynamics. In order to investigate this we define three cell boundary descriptions of varying complexities for each of three widely used off-lattice individual-based models: overlapping spheres, Voronoi tessellation, and vertex models. We apply our models to multiple biological scenarios to investigate how cell boundary description can influence tissue-scale behaviour. We find that the Voronoi tessellation model is most sensitive to changes in the cell boundary description with basic models being inappropriate in many cases. The timescale of tissue evolution when using an overlapping spheres model is coupled to the boundary description. The vertex model is demonstrated to be the most stable to changes in boundary description, though still exhibits timescale sensitivity. When using individual-based computational models one should carefully consider how cell boundaries are defined. To inform future work, we provide an exploration of common individual-based models and cell boundary descriptions in frequently studied biological scenarios and discuss their benefits and disadvantages.

## 1 Introduction

Epithelial tissues line bodily surfaces including the skin (Nafisi and Maibach, 2018), intestines (Reed and Wickham, 2009), respiratory tract (Hermans and Bernard, 1999), blood vessels (Stolz and Sims-Lucas, 2015), cornea (Frost et al., 2014) and sweat glands (Sundberg et al., 2018). The primary functions of epithelial tissues are to protect organs, secrete enzymes and hormones and absorb harmful substances. As epithelial tissues are more exposed to external impacts than other tissue types, cancer develops more frequently in epithelial tissues (Hinck and Näthke, 2014). Epithelial cancers are known as carcinomas and can spread quickly (Becker et al., 2017; Hsieh et al., 2017). Certain forms of carcinoma, such as high-grade serous ovarian cancer and lung adenocarcinoma, severely impact patients’ quality of life and have low survival rates (Chandra et al., 2019; Lisio et al., 2019; Lu et al., 2019). In addition to the increased risk of cancer, the exposure of epithelial tissues to the external environment may also lead to the tissue being damaged in non-cancerous settings. Disruption in epithelia due to injury can leave organs vulnerable to infection, or prevent them from functioning normally (Evans et al., 2013; Croasdell Lucchini et al., 2021; Subramanian et al., 2020). Given their ubiquity throughout the body and importance in maintaining organ health, epithelial tissues have been extensively studied *in vitro, in vivo* and *in silico*. However, there remain many unknowns about epithelial behaviour, including which cellular processes and mechanisms are involved in epithelial homeostasis and morphogenesis (Kaliman et al., 2021; Kondo and Hayashi, 2015).

Individual-based modelling is a popular tool for simulating biological phenomena in epithelial tissues such as wound healing, tumour growth and colorectal cancer (Osborne et al., 2017). Epithelial tissues generally consist of confluent layers of cells. The polygonal structure and confluent arrangement of cells in epithelial tissues lend themselves well to individual-based modelling. Alternatively, epithelial tissues can be modelled as a continuous chain, sheet, or block of interacting cells, using techniques within continuum mechanics. An advantage of individual-based models, compared to their continuum tissue-scale counterparts, is that they are able to explicitly incorporate individual cell behaviour, such as cell cycles, cell interactions, and cell phenotype proportions (Osborne et al., 2017). The first individual-based models were on-lattice models, usually of cell-sorting, where cell locations were restricted to lattice sites (Brodland, 2004). On-lattice models are computationally less expensive and conceptually more straightforward than their more recently developed off-lattice, or lattice-free, counterparts (Van Liedekerke et al., 2015). Historically, the use of individual-based models, particularly off-lattice models, have been limited by computational complexity. However, as both computational costs decrease and computational power increases, individual-based models are increasing in usage and detail (Fig. 1). Furthermore, as additional model functionality has developed, individual-based models can also describe subcellular behaviour, such as cell responses to signalling pathways (Sun et al., 2009; Buske et al., 2011; Germano and Osborne, 2021; Sandersius and Newman, 2008). These tools allow for a range of biological scenarios that were previously inaccessible to the computational biologist to be explored.

**Fig. 1.**
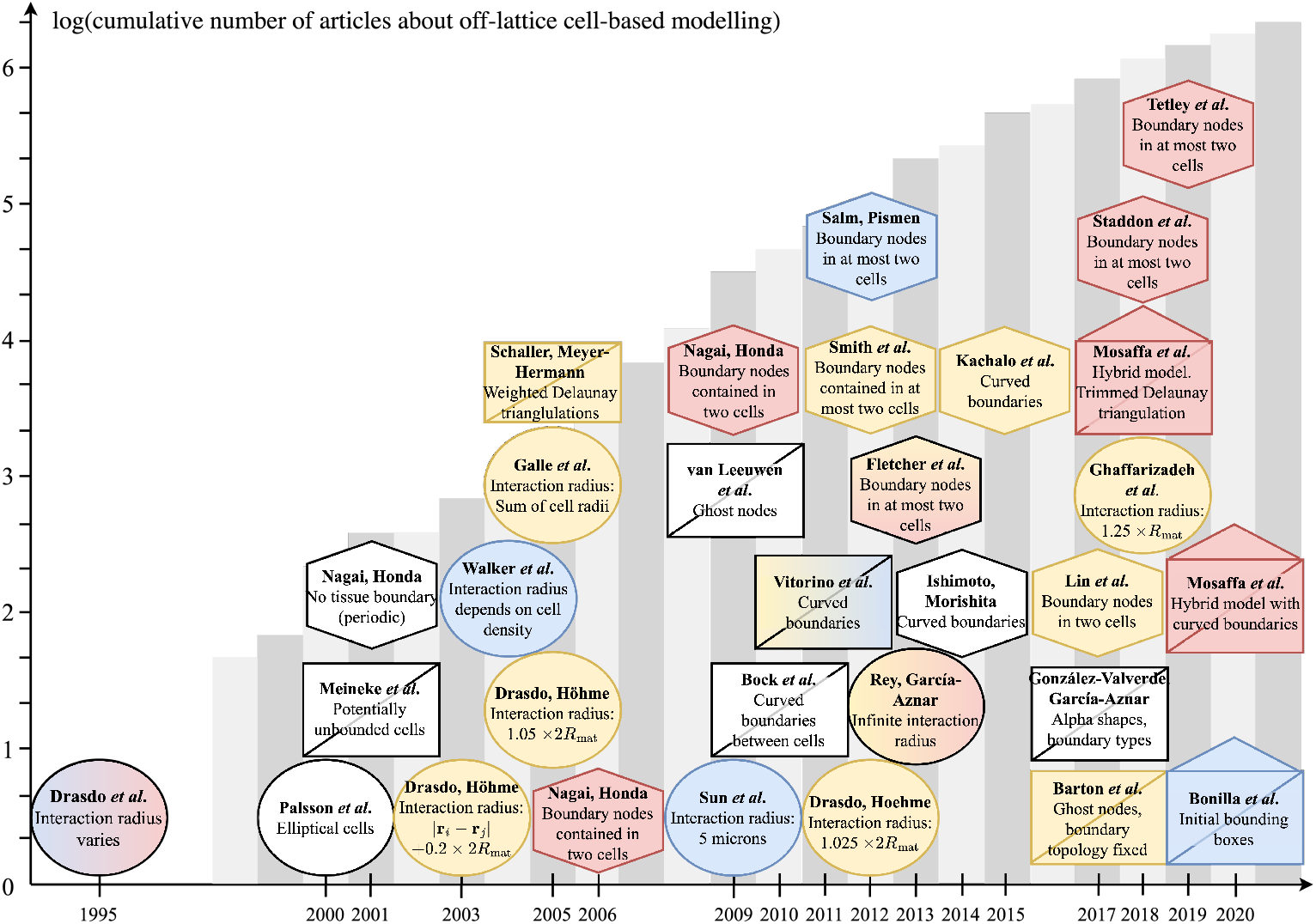
Timeline of individual-based tissue models (Drasdo et al., 1995; Palsson and Othmer, 2000; Meineke et al., 2001; Nagai and Honda, 2001; Drasdo and Höhme, 2003, 2005; Galle et al., 2005; Schaller and Meyer-Hermann, 2005; Nagai and Honda, 2006; Sun et al., 2009; Van Leeuwen et al., 2009; Nagai and Honda, 2009; Bock et al., 2010; Vitorino et al., 2011; Smith et al., 2012; Drasdo and Hoehme, 2012; Salm and Pismen, 2012; Rey and Garcia-Aznar, 2013; Fletcher et al., 2013; Ishimoto and Morishita, 2014; Kachalo et al., 2015; Barton et al., 2017; González-Valverde and García-Aznar, 2017; Lin et al., 2017; Ghaffarizadeh et al., 2018; Mosaffa et al., 2018; Staddon et al., 2018; Tetley et al., 2019; Mosaffa et al., 2020; Bonilla et al., 2020). Author names are in bold text, with a description of the boundary type used in plain text. The centre of the shape corresponds to the year of publication along the horizontal axis. Circles represent overlapping spheres models, rectangles with a diagonal line represent Voronoi tessellation models and hexagons represent vertex models. Rectangles with a triangle on top represent models that are hybrid models (using forces on both cell centres and polygonal vertices). Red coloured shapes have void closure applications with little or no cell proliferation, yellow shapes have tumour growth applications and blue shapes represent colliding tissue front applications. Shapes that have multiple colours are papers with multiple applications. White coloured shapes have other applications, but are significant for their contributions on the discussions of cell boundaries. The height of the grey bars in the background represent the natural logarithm of the cumulative number of papers up to and including the corresponding year that can be found using the search terms ‘off-lattice individual-cell model’ using Google Scholar. Note that this terminology first appears in the literature in 2003, therefore the bars corresponding to the years from 1997 to 2004 show the natural logarithm of the cumulative number of citations of early publications of each model type: OS (Drasdo et al., 1995), VT (Meineke et al., 2001) and VM (Nagai and Honda, 2001)

In this work, we focus on off-lattice models. These models are more realistic and are based on physical principles. Moreover off-lattice models are likely to continue to increase in popularity into the future, as they provide a rigorous framework for investigating mechanistic behaviour. In off-lattice models, cells are free to move in space and are not restricted to lattice sites (Brodland, 2004). Off-lattice models can be further categorised into cell-centre models (where a cell is defined by a single point in space) and element based models (where a cell is defined by more than one point in space). One of the computationally-simplest cell-centre models is the overlapping spheres (OS) model (Drasdo et al., 1995). In this model, cells are modelled as point-particles in space, which represent the cell. Cell centres will interact with each other if they are within a certain distance of each other. If they are too close, there is a repulsive force between the cell centres. Whereas if the distance between the cell centres is within a specified range, the cell centres experience an attractive force. An alternative cell-centre approach is known as a Voronoi tessellation (VT) model (Meineke et al., 2001). In this model, whether cells interact with each other is determined by the Delaunay triangulation of the cell centres (Boots et al., 2009), where the Delaunay triangulation of a set of points partitions the space into triangles (in ℝ^2^), or tetrahedrons (in ℝ^3^) such that no point lies in any circumcircle/sphere of the triangles/tetrahedrons. If there is an edge/face between two cell centres in the Delaunay triangulation, then the cell centres exert forces on each other. Typically the form of this interaction is a simple linear spring acting between the cell centres (Pathmanathan et al., 2009). The dual of the Delaunay triangulation is the Voronoi tessellation (the space is partitioned into polygons such that all points within a particular polygon are closer to the enclosed cell centre compared to any other), which is used to describe the shapes of the cells (Boots et al., 2009). Cell-centre models are considered to be a zeroth order approximation to the physical system, as they typically model all mechanical intra- and extra-cellular interactions with a single mechanism.

An alternative approach to off-lattice modelling that does not use cell centres is the vertex model (VM) (Nagai and Honda, 2001). In this model, cells are represented by polygons, with forces acting on the vertices of the polygons. These forces arise due to cell compressibility and cytoskeletal tension and adhesion between cells. As the vertices move and cells undergo death or division, rearrangements are required to maintain the structure of the tissue (Nagai and Honda, 2001, 2006). Within a VM, cell compressibility, cytoskeletal tension and adhesion between cells each has its own mathematical mechanism of delivery; the VM is considered a first order approximation to the physical system.

Although there have been several published works using off-lattice models to investigate various biological problems in tissues (Fig. 1), there has been no investigation to date into the influence of the choice of cell boundary descriptions for these models. Here we consider the effects of cell boundaries in two-dimensional models. An extension to three dimensions may result in additional findings and be of further use when three-dimensional models are computationally cheaper to perform. However, as three-dimensional models are not widespread, here we focus on two dimensions. There have been multiple studies into more complicated cell boundaries than those we explore here. Depending on the context, more complex models may be useful, such as immersed boundary and finite element models (Rejniak, 2007; Chen and Brodland, 2000). However, these more complex models are generally computationally intensive. As such, these models are limited in the number of cells that can be feasibly simulated. We limit our scope to three different cell boundary descriptions per model. These three boundaries are canonical examples of boundaries of varying complexity. We seek to highlight the influence, if any, of having increasingly complex cell boundaries on epithelial tissue dynamics. In this work we explore commonly used cell boundary descriptions for the OS and VT models and VM to determine their impact on the results of simulations for three biological scenarios: void closure, tissue growth and tissues colliding, shown in Fig. 2.

**Fig. 2.**
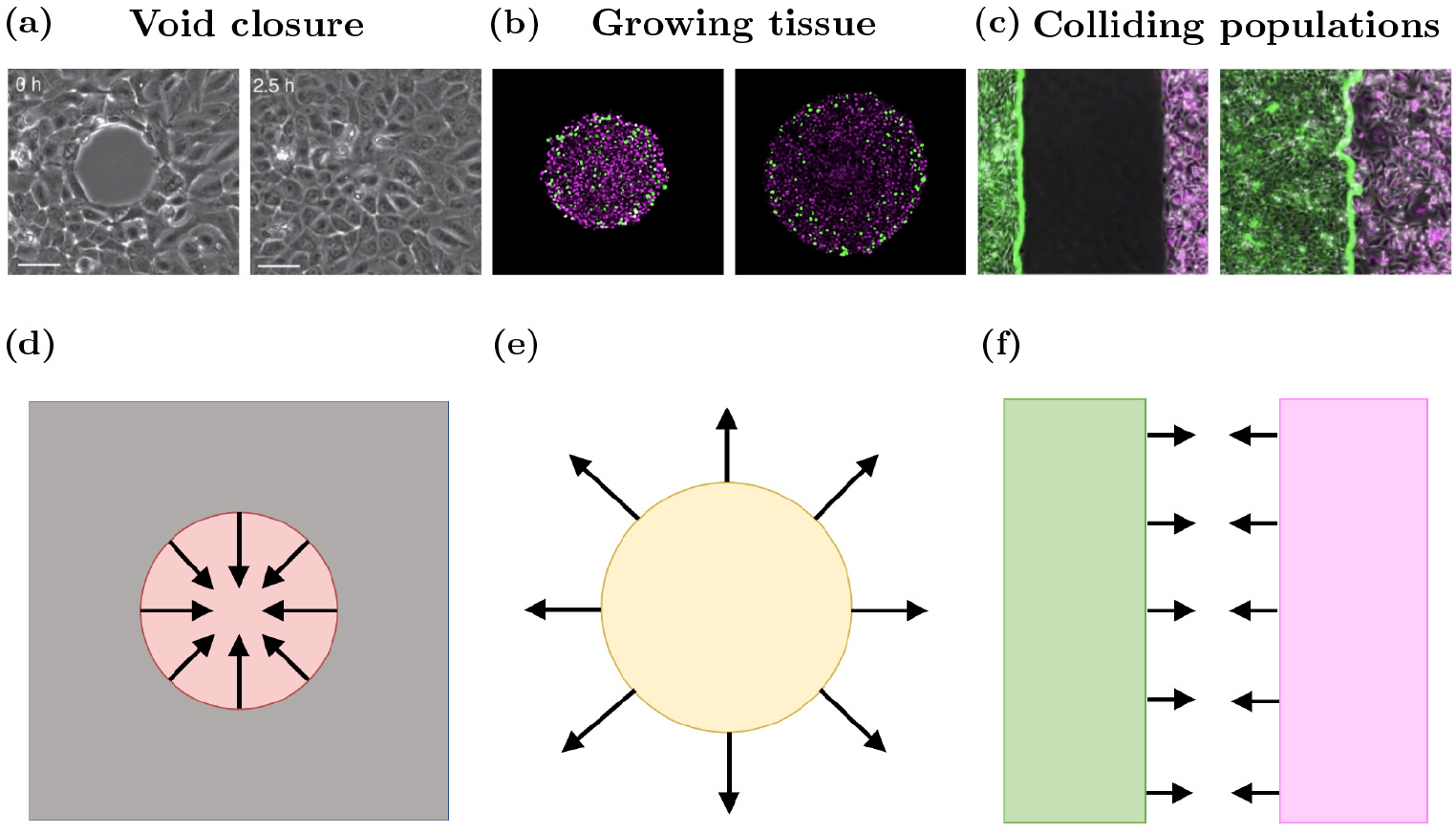
Model schematics for void closure, tissue growth and tissue collisions. *In vitro* snapshots of (a) wound closure (b) tumour growth, and (c) colliding tissues. Left panels of *in vitro* figures are early time, right panels are later times. Model schematics of (d) void closure, (e) a growing tissue and (f) tissue collision. Images are adapted from (Vedula et al., 2015), (Murphy et al., 2022) and (Heinrich et al., 2022)

The remainder of the paper is structured as follows. Firstly, we provide an overview of each of the three model types considered in this work and introduce the different cell boundary descriptions investigated for each model. Next, we explore each biological scenario using the different model types and boundary descriptions and present the results as separate case studies. Finally we use our simulations to inform our assessment of which of the nine (3 off-lattice models *×* 3 cell boundary descriptions) models are most appropriate for each biological scenario, and discuss the benefits and drawbacks of each model and boundary description.

## 2 Methods

We begin this section by describing the cell-centre models -OS and VT - and the VM. Further details of each model can be found in the literature (Osborne et al., 2017). After introducing the models, we define three cell boundary descriptions per model to be explored, from simplest to most complex. A schematic of the different boundary descriptions for the three models investigated is given in Fig. 3. A timeline of the development of each model, and which boundary descriptions have been used in previous studies is given in Fig. 1.

**Fig. 3.**
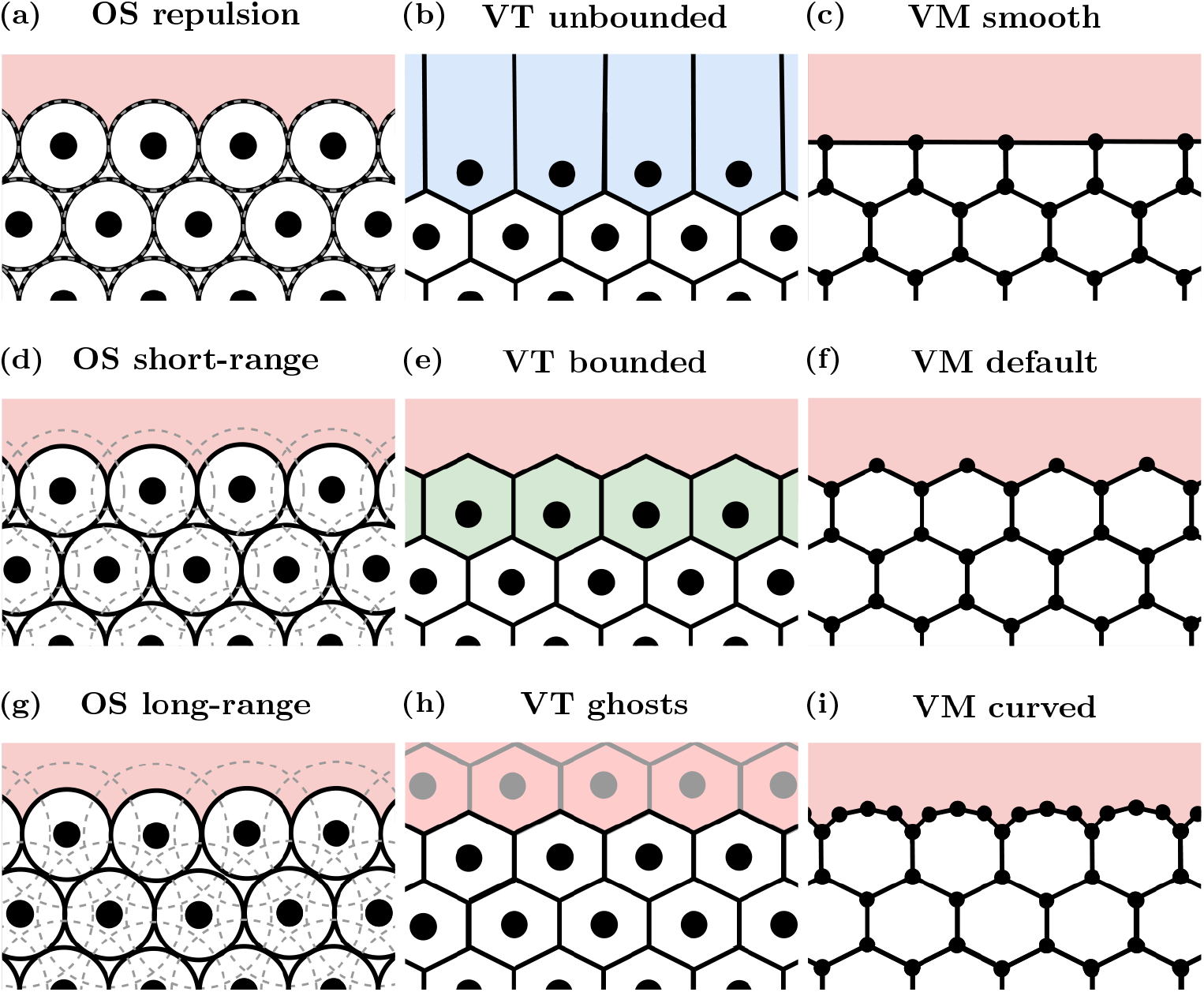
Cell boundary description schematics. OS models are pictured in (a), (d) and (g) (left column); VT models are shown in (b), (e) and (h) (middle column); and VMs are shown in (c), (f) and (i) (right column). The most computationally efficient boundaries are shown in (a)-(c) (top row). The most commonly used boundaries are displayed in (d)-(f) (middle row). The most computationally complex boundaries are shown in (g)-(i) (bottom row)

### 2.1 Cell-centre models

We first define the cell-centre models (OS and VT). In cell-centre models we represent cells by a point, free to move in space, that represents the centre of the cell. Suppose that there are *N* (*t*) cells at time *t*. Let **r**_*i*_ (*t*) *=* (*x*_*i*_ (*t*), *y*_*i*_ (*t*)) *∈* **ℝ**^2^ denote the position of the centre of cell *i ∈* {1,…, *N* (*t*)}. Note that these methods are generalisable to ℝ^*n*^, for *n* ∈ {1, 2, 3}, however we restrict our focus to two-dimensional models in this work. We make the simplifying assumption that all cells exhibit the same mechanical properties. Subsequently, force balances are used to derive equations of motion for the cell centres. We assume that cells are moving through dissipative environments where the inertial forces are small compared to dissipative forces (Dallon and Othmer, 2004). Therefore, the equation of motion for cell *i* in the cell centre models is given by:

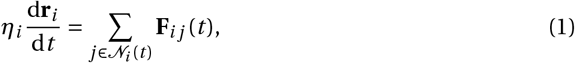

where *η*_*i*_ is the drag coefficient, **F**_*i j*_ (*t*) is the interaction force acting between cell *i* and its neighbours *j* ∈ 𝒩_*i*_ (*t*), where *𝒩*_*i*_ (*t*) denotes the set of all neighbours of cell *i* at time *t*. To solve the equation of motion numerically, the forward Euler method is used:

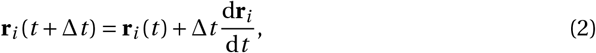

where ∆*t* is the time step to be used, chosen to ensure numerical stability.

To allow tissues to grow, we introduce a mechanism for cells to divide. Specifically, a Bernoulli trial in conjunction with contact inhibition is used to determine whether or not a cell divides, thereby introducing stochasticity into the simulations. If a cell is more than *t*_div_ hours old, the probability of the cell dividing within the subsequent hour is *p*_div_. However, the cell must have an area greater than or equal to *A*_*q*_ to be able to divide. If a cell’s area is less than *A*_*q*_, then the probability of the cell dividing is zero, and the cell is considered *quiescent*. If the cell grows beyond a size of *A*_*q*_, then it is able to divide again. Cells divide symmetrically, meaning their offspring have the same proliferative properties. This is a common model of malignant cell division, where the only thing preventing a cell dividing is space constraints. When a cell divides the cell centre of the daughter cell is placed a small distance from the parent cell’s cell centre in a randomly selected direction.

#### 2.1.1 Overlapping spheres

For the OS model, we define the force acting between cells *i* and *j*, **F**_*i j*_ (*t*), as (Osborne et al., 2017):

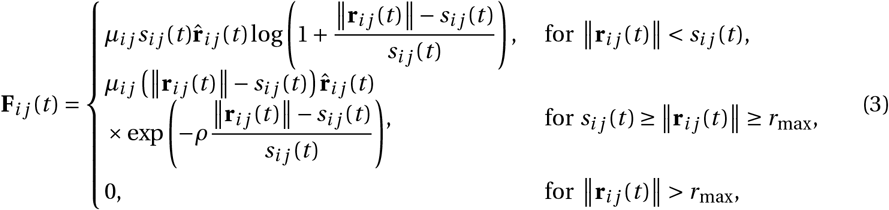

where *µ*_*i j*_ *= µ* is the magnitude of elastic interaction between cells *i* and *j* and *s*_*i j*_ (*t*) is the natural separation between cells *i* and *j*, given by *s*_*i j*_ (*t*) *= R*_*i*_ (*t*)*+R*_*j*_ (*t*). Here, *R*_*i*_ (*t*) is the radius of cell *i*, and is constant for mature cells. To avoid sudden changes in the potential of the tissue and for consistency with existing models, during the first hour after cell division, *R*_*i*_ (*t*) increases linearly from half of the mature radius to the mature radius (Osborne et al., 2017). For simplicity, we assume the mature cell radius is the same for all cells, denoted *R*_mat_. The vector from cell *i* to cell *j* at time *t* is denoted **r**_*i j*_ (*t*) *=* **r**_*j*_ (*t*)*−***r**_*i*_ (*t*), with unit vector 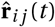. The parameter *ρ* determines the decay in the attractive force between two cells. We note that for ║**r**_*i j*_ (*t*)║ *< s*_*i j*_ (*t*), cells repel each other, and for *s*_*i j*_ (*t*) *≥* ║**r**_*i j*_ (*t*)║ *≥ r*_max_, they attract each other.

In the OS model, cells are point particles, with no volume or ‘boundary’. However, the interaction forces (and therefore interaction radius *r*_max_) between cells implicitly act as a boundary condition as they affect which cells are considered to be neighbours (and therefore defines 𝒩_*i*_ (*t*)). Here we investigate three different interaction radii for the OS model to simulate different cell boundary descriptions. These interaction radii represent:

- **Repulsion**: In these simulations, the interaction radius is 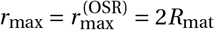, so there is no attraction between neighbouring mature cells (Fig. 3(a)). However, there will be attraction between immature cells (cells with an age less than one hour) if the distance between their cell centres is less than 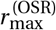 and greater than the sum of their current
- **Short-range**: Here the interaction radius is 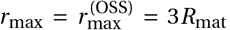, representing both repulsion and some attraction between mature neighbours, as shown in Fig. 3(d). When the distance between two cell centres of mature cells is less than 2*R*_mat_, the cells exert repulsive forces on each other. Whereas when the distance between the cell centres is between 2*R*_mat_ and 3*R*_mat_, the cells exert attractive forces on each other.
- **Long-range**: In this regime the interaction radius is 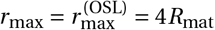. The region of attraction between neighbours is larger in this setup than the short-range case, as can be seen in Fig. 3(g). As in the short-range case, when the distance between two cell centres of mature cells is less than 2*R*_mat_, the cells exert repulsive forces on each other. Whereas when the distance between the cell centres is between 2*R*_mat_ and 4*R*_mat_, the cells exert attractive forces on each other.

The lack of volume for cells in the OS model presents other challenges, as cell cycle descriptions or subcellular mechanics may rely on cell volume to determine whether or not a cell should divide or exert additional forces on its neighbours. In these scenarios, approximations for cell volume are employed. For the purposes of this investigation, the area of cell *i, A*_*i*_ (*t*), is defined as (Osborne et al., 2017):

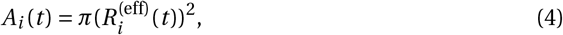

Where

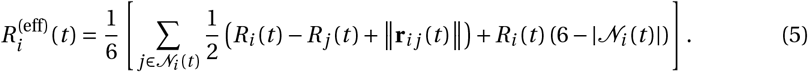

#### 2.1.2 Voronoi tessellation

In the VT model, which cells are neighbours is determined by the Delaunay triangulation of cell centres. Recall that the Delaunay triangulation of a set of points in ℝ^2^ partitions the space into triangles such that no point lies in any circumcircle of the triangles. If there is an edge between two cell centres in the Delaunay triangulation, then the cell centres exert forces on each other. The force acting between cell *i* and its neighbours *j ∈ 𝒩*_*i*_ (*t*) in the triangulation is given by (Pathmanathan et al., 2009; Meineke et al., 2001):

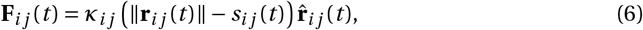

where *κ*_*i j*_ *= κ* is the linear spring constant, and *s*_*i j*_ (*t*) is the natural separation between cells *i* and *j*. In the VT model, the natural separation between neighbouring cell centres increases linearly from 0.1 to 1 cell diameter during the first hour after cell division, preventing abrupt changes in the potential of the tissue and consistent with existing models (Osborne et al., 2017).

Similar to the OS model, when we discuss cell boundaries, we focus on which cells interact with each other. Unlike OS models, VT models include explicit descriptions of cell shapes, and therefore the concept of cell area is mostly well-defined. However, the standard VT tessellates over the entirety of ℝ^2^, allowing cells on the outer boundaries of the tissue to have infinite area. Furthermore, since there is no limit on how long an edge in the Delaunay triangulation may be, connections may be formed between cell centres that are far apart, causing cells to interact in ways that are biologically infeasible. An approach for avoiding this issue is to use a bounding polygon. All edges extending beyond the polygon are shortened so their end point lies on the polygon. This also bounds the Delaunay triangulation, removing the unrealistic interactions between distant cells. An alternative method for bounding a tessellation in the context of cell modelling is to introduce *ghost nodes*. Ghost nodes are nodes that surround the boundary of a tessellation, but are not a part of the biological problem being considered. Ghost nodes themselves may have infinite area, but the ‘real’ nodes that describe the cells will have finite area. The three types of VT cell boundary descriptions considered in this work are:

- **Unbounded VT**: In these simulations, the VT (and Delaunay triangulation) is used as per the standard definition, meaning that cells at the free boundaries of the tissue may have infinite area (in practice, the area is bounded by numeric limits). Moreover, edges in the Delaunay triangulation may be long, leading to unrealistic interactions between cell centres. A schematic for this model is given in Fig. 3(b).
- **Bounded VT**: Under this model, at every time step, additional nodes are added around the boundary so the tessellation defining the tissue has finite volume (the cells on the boundary have finite area), as shown in Fig. 3(e). Note that the additional nodes used here are only present during the tessellation calculation, and are destroyed every time step after the tessellation is created. The placement of these nodes and subsequent VT is described in Appendix A. We introduce a cut-off length, 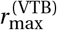, in the Delaunay triangulation, analogous to the OS model: if an edge in the triangulation is greater than 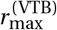, then it is removed and the cells connected by that edge do not interact.
- **Ghost nodes**: Ghost nodes are additional nodes beyond the tissue boundary, similar to the ones introduced in the bounded VT model, except that they persist over time and react to forces exerted by the ‘real’ nodes (nodes representing cells in the simulation). In practice, ghost nodes are created by generating more cells than are necessary and labelling all of the cells outside of the representation of the tissue as ‘ghosts’. A schematic is provided in Fig. 3(h). Ghost nodes exert forces on each other, but do not exert forces on the real nodes in the tissue.

For the void example, ghost nodes will be removed if their associated Voronoi area is below a certain threshold, 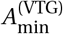. In the colliding tissues example, ghost nodes are also removed if the associated Voronoi area is less than 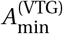 or if they are not connected to any other ghost nodes.

### 2.2 Vertex model

Unlike the OS and VT models, the VM does not include forces acting on cell centres. Instead, cells are modelled as non-overlapping polygons with forces acting on the vertices of the polygons. In the VM, we consider the points {**r**_1_(*t*), …, **r**_*N*_vert_(*t*)_} (here, the set of vertices is greater than the number of cells in the system). The force on each vertex is derived from an energy function, a viscous drag term, and again involves an assumption that cells are moving through highly viscous environments where the inertial forces can be ignored (Dallon and Othmer, 2004). Balancing these forces leads to an equation of motion for vertex *i* (Fletcher et al., 2013):

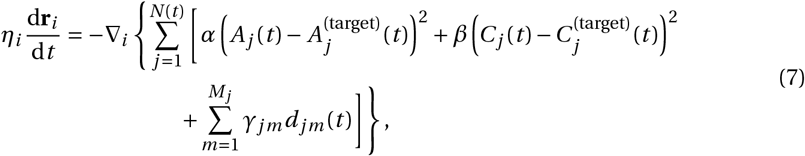

where *α* and *β* are positive constants, describing cell compressibility and cytoskeletal tension, respectively. We define *A*_*j*_ (*t*) and *C*_*j*_ (*t*) as the area and perimeter of cell *j* at time *t*, respectively. Each cell is prescribed a target area, with the target area of cell *j* at time *t* denoted by 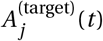. For simplicity, we assume that every mature cell has the same target area, *A*_target_. The target area for mature cells is constant. During the first hour after cell division, the target area increases linearly from 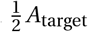 to *A*_target_, preventing abrupt changes in the potential of the tissue and consistent with existing models (Fletcher et al., 2013; Kursawe et a 2015). Each cell also has a target perimeter. The target perimeter of cell 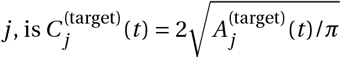. The number of vertices contained in cell *j* is denoted by *M*_*j*_. We assume that the vertices of each cell are ordered in an anti-clockwise fashion. The parameter *γ*_*jm*_ is a positive constant describing cell adhesion, and depends on whether the edge connecting vertices *m* and *m +* 1 (i.e. the edge between vertex *m* and the next vertex in anti-clockwise order) in cell *j* is an edge on the boundary, an edge shared between two cells of the same type or an edge shared between two cells of different types. The length of the edge connecting vertices *m* and *m +* 1 in cell *j* at time *t* is *d*_*jm*_ (*t*).

To ensure that cells do not overlap or deform in unrealistic ways, rearrangements of the cells and vertices may be necessary (Fletcher et al., 2013). Suppose that an edge connects vertices between cells *A* and *B*, who share no vertices, and that cells *C* and *D* are adjacent to the edge. If the edge has length less than the rearrangement threshold, *d*_*r*_, then the edge is replaced with a perpendicular longer edge such that cells *C* and *D* are no longer adjacent (they share no vertices), but cells *A* and *B* become adjacent. If a cell area is too small, below some threshold *A*_min_, then the cell is removed from the simulation. Finally, if a cell intersects another cell (or cells), then nodes may be inserted or replaced. Existing operations for mesh restructuring are described in more detail in (Fletcher et al., 2013). Further cases are developed as necessary for this work, and can be found in Appendix B. The cell boundary descriptions used in the VM are:

- **Smooth boundaries**: We define a smooth boundary vertex simulation as one in which a vertex is shared by at least two cells. Consequently, any vertices on the tissue boundaries contained in only one cell are removed. This has the effect of ‘flattening’ cells at the boundaries, as seen in Fig. 3(c).
- **Default boundaries**: In existing VMs, nodes are typically initialised using a (finite) VT of cell centres that are prescribed by the user. These nodes then move over time according to the force laws described by the equations of motion in Equation (7). This usually results in boundary cells that contain one node that is contained in only one cell, two nodes contained in two cells and the remaining nodes are internal nodes that are contained in three cells, as in Fig. 3(f). By neither adding nor removing additional nodes along the boundary edges, the shapes of the boundary cells are less constrained than the smooth boundary case.
- **Curved boundaries**: A curved boundary vertex simulation is where additional nodes are added to boundary cells to approximate curved cell boundaries, as shown in Fig. 3(i). To implement this, we define a maximum boundary edge length, 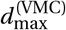, for boundary edges (note that this does not apply to internal edges). If a boundary edge length exceeds the maximum edge length, an additional node is inserted at the midpoint along the edge. We also define a minimum boundary edge length, 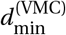, for boundary edges, which acts in a similar manner. If an edge length is less than the minimum edge length, the two nodes connected by the edge are removed and replaced by a node in the midpoint of the edge.

### 2.3 Implementation

All simulations are performed in the open source simulation package Chaste (Mirams et al., 2013; Pitt-Francis et al., 2009; Cooper et al., 2020). Spatial units for the simulations are non-dimensionalised to cell diameters (CDs) and cell mass (CM), and time units are in hours. Parameter values used across simulations are provided in Table 1. The code used to produce this work is available at https://github.com/jmosborne/TissueBoundaries.

**Table 1.**
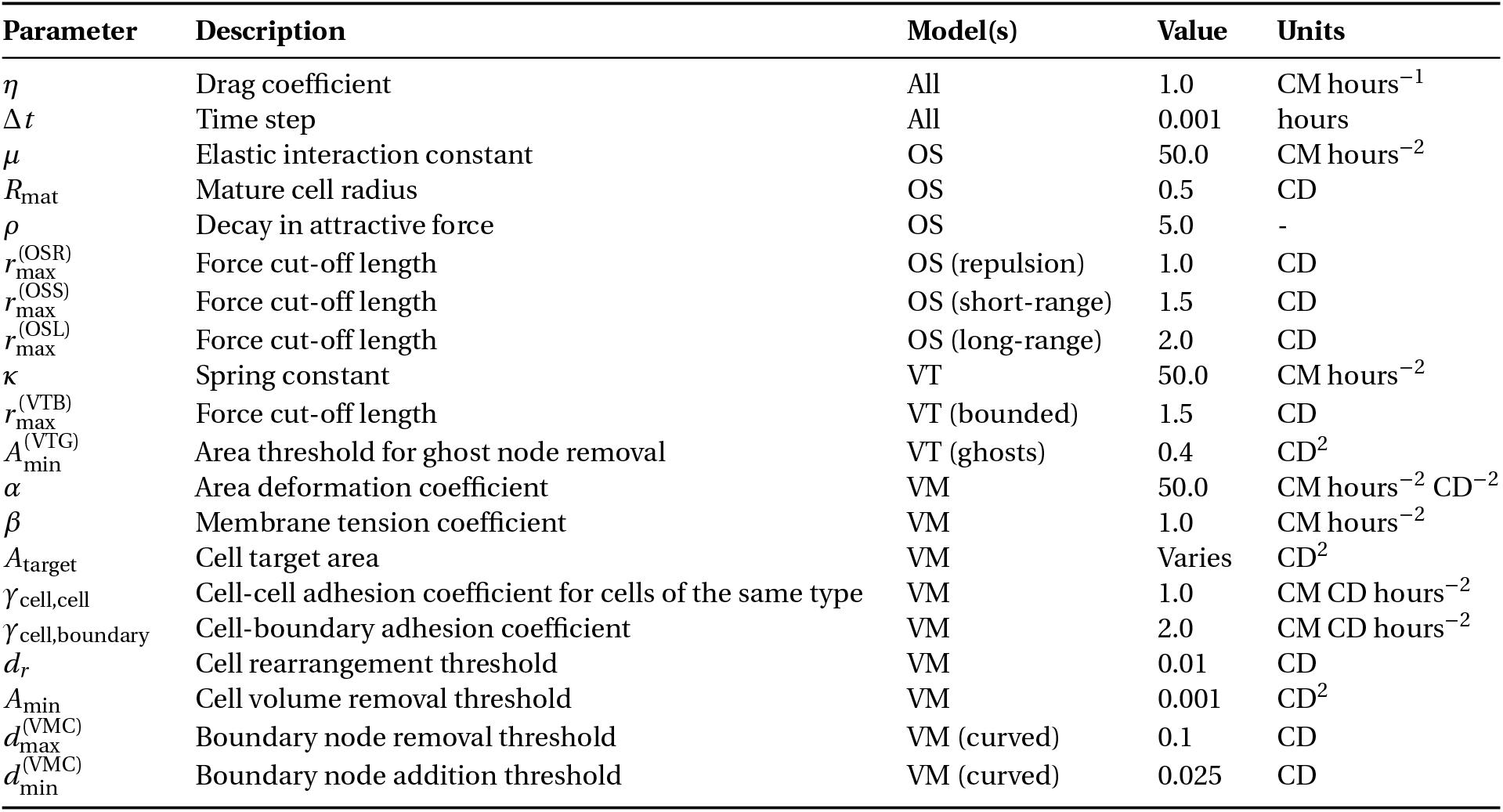
Parameter values used across all simulations. Parameter values sourced from (Pathmanathan et al., 2009; Fletcher et al., 2013), except for boundary specific values, rearrangement thresholds and adhesion between cells and free space in the VM

**Table 2.** Model comparisons for void closure.

|  |  | Model |  |  |
| --- | --- | --- | --- | --- |
|  |  | OS | VT | VM |
|  | **Low** | **Repulsion:** Quickest closure time (of OS models), tissue tearing occurs. | **Unbounded:** Void not possible, cells where void would be are large, but decrease in size over time (of VT models). | **Smooth:** Quickest closure time (of VM models), homogeneous cell sizes after void closure. |
| **Boundary resolution** | **Default** | **Short-range:** Quick closure time (OS), heterogeneous cell sizes after void closure. | **Bounded:** Jump discontinuities in area and changes in void shape due to edge changes in Delaunay triangulation. | **Default:** Quick closure time (VM), homogeneous cell sizes after void closure. |
|  | **High** | **Long-range** Slowest closure time (OS), after void closure larger cells in the space where void was surrounded by smaller cells. | **Ghost nodes:** Jump discontinuities in area and changes in void shape due to ghost node removal. | **Curved:** Slowest closure time (VM), homogeneous cell sizes after void closure. |

## 3 Results

In this section we describe the differences between model type arising from the choice of cell boundary descriptions in each of our biological scenarios. We find that tissue behaviour often differs between model types, and at a more granular level, there is also significant variability between cell boundary descriptions.

### 3.1 Void closure: Choice of model and representation dictates void closure timescale and mechanism

The first biological scenario we consider is an internal void in a tissue, such as a wound or the final stages of morphogenesis (Brugués et al., 2014; Vedula et al., 2015; Tetley et al., 2019; Ajeti et al., 2019) Biological experiments into void closure have attempted to determine how cytoskeletal structures impact void closure (Brugués et al., 2014) (Fig. 2(a)). However, the mechanisms driving void closure are yet to be fully understood. In these biological contexts, the nature of cell-cell and cell-environment interactions and collective cell behaviours are not fully understood. For example, there have been multiple mechanisms of epidermal cell migration during wound healing that depend on cell-cell and cell-environment interactions (Rousselle et al., 2019). However, it is yet to be determined which mechanisms are applicable in different contexts. Individual-based models are useful for exploring hypotheses about cell scale behaviour. Examples of individual-based models of void closure are shown in red in Fig. 1. However, artefacts introduced by the choice of individual-based model and cell boundary description can impact the characteristics of void closure.

A common metric of interest in this scenario is the time to void closure. The time to void closure may be impacted by cell behaviour, specifically, which cells interact with each other and how. Computational individual-based models are able to explicitly incorporate inter-cellular interactions. Furthermore, unlike continuum models, the shape of the void does not significantly impact the complexity or fundamental structure of individual-based models, making them an ideal framework for exploring the evolution of void shape during closure. In this section we consider a simplified void closure scenario, where tissue compression leads to void closure, and investigate how different cell boundary descriptions change both the time to void closure and the wound geometry. *In vitro* images of wound closure and a model schematic are shown in Fig. 2(a) and (d).

In 2009, Nagai and Honda used their VM to investigate wound closure driven by tissue compression *in silico* (Nagai and Honda, 2006, 2009). Here we use a similar approach to understand what effects, if any, model choice has on compression-driven void closure. We begin by initialising a hexagonally-packed tissue 14 cells wide and 16 cells high. We choose hexagonal packing so any cell movement is not caused by the tissue relaxing to the least energetic configuration. We implement a periodic domain for cell motion in both the *x* and *y* -directions to simulate a much larger tissue than would otherwise be computationally feasible. This tissue is then compressed in both the *x* and *y* -directions to a length that would be optimally packed for a tissue 12 cells wide and 13 cells high and allowed to relax to its equilibrium. Cells are subsequently removed from the tissue if their cell centre resides within a polygonal (capsule) region defined by |*y − x*| *<* 2, with 4 *< x, y <* 8, where *x* and *y* are the *x* and *y* -coordinates of the cell centre, creating a void in the tissue. The tissue is now allowed to relax to in-fill the space. Cells do not divide in these simulations, and so the simulations are deterministic. Traces of the void over time for each model are given in Fig. 4 and void area over time is shown in Fig. 5. Calculations of void area are described in Appendix C. Void closure is considered to have occurred when the void area reaches 0 CD^2^.

**Fig. 4.**
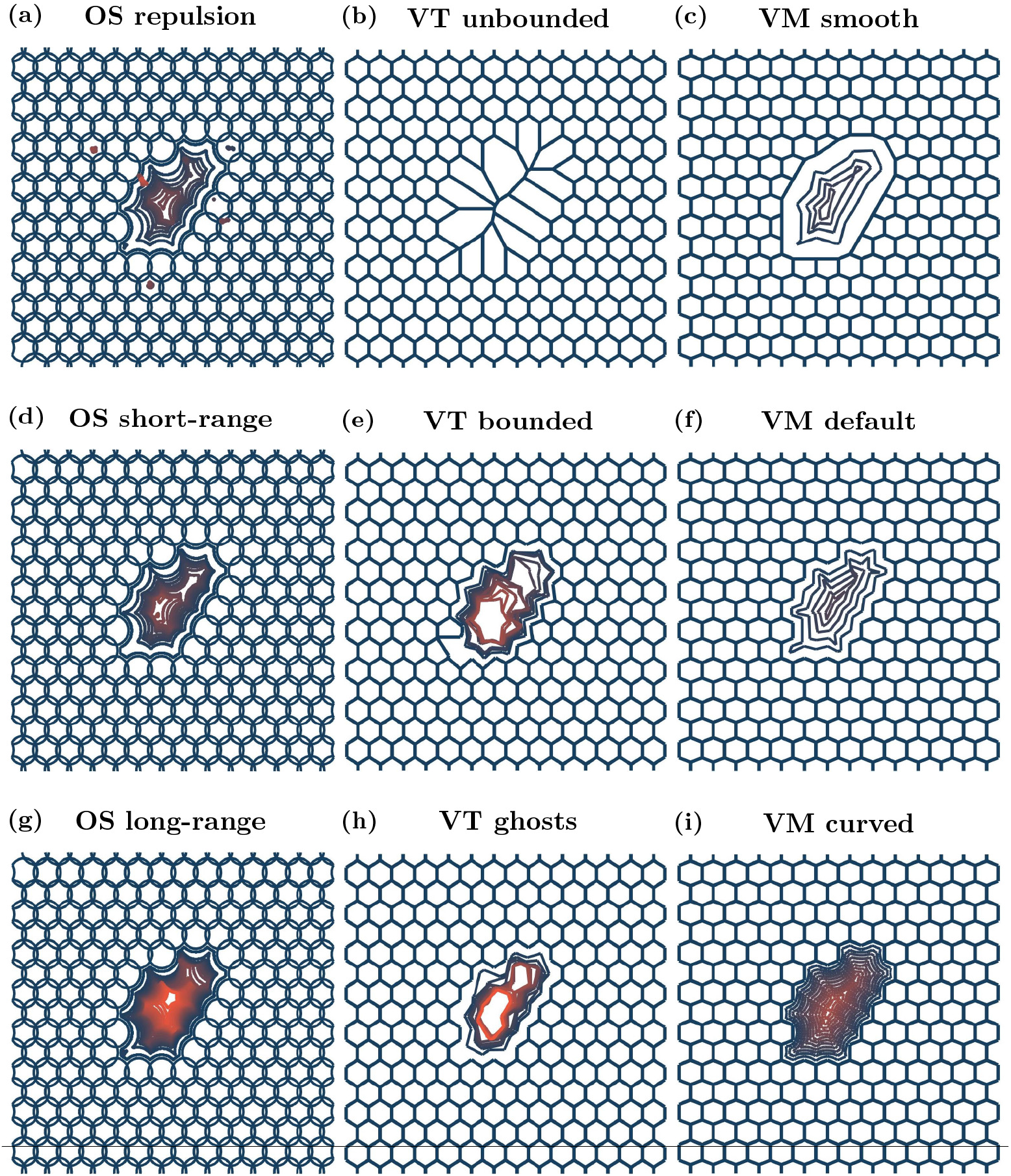
Void outlines over time. Initial cell positions and configuration in blue, and void boundary trace for each model setup at later times. Orange outlines correspond to void boundary traces at later times, if the void has not closed. Void outlines are plotted every 0.04 hours until the void has zero area or 20 outlines have been plotted, whichever occurs first. The small ‘dots’ seen in (a), the OS repulsion case, are small tears that occur in the tissue at various times due to how the boundaries are defined, see Appendix C. Videos of all simulations are provided in the Supplementary Video 1

**Fig. 5.**
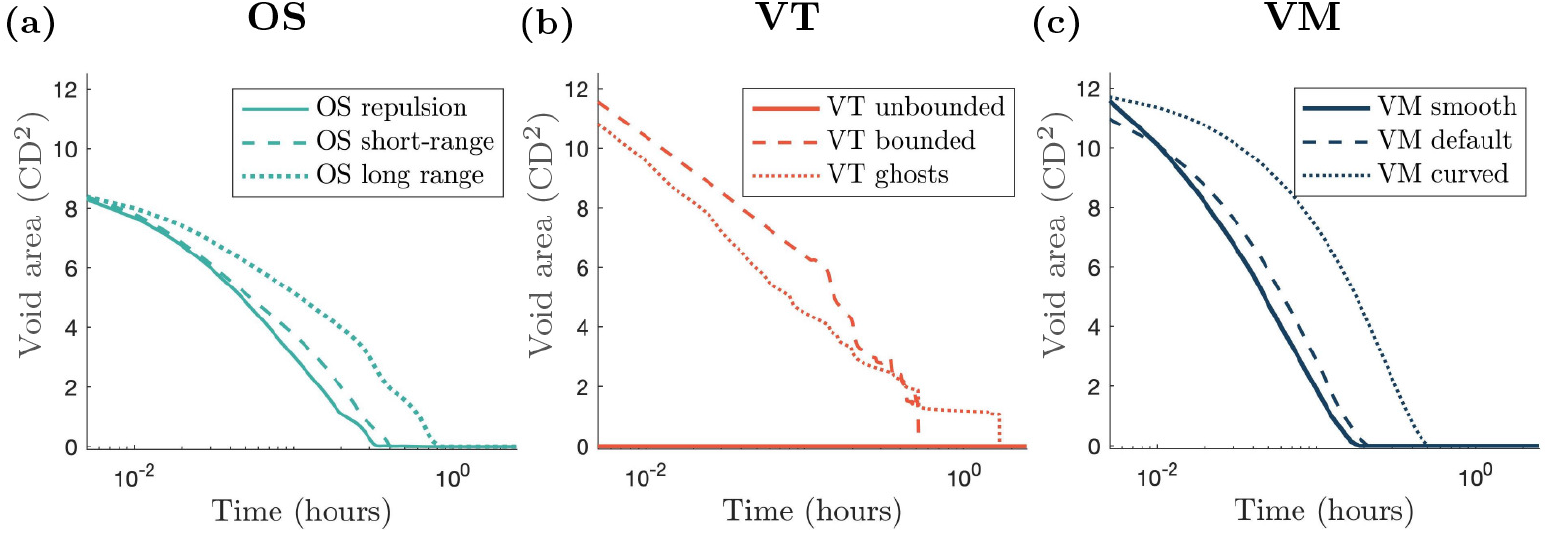
Void area over time curves for each model. The curve corresponding to the simplest cell boundary description for each model is plotted using a solid line and the curves corresponding to the most computationally complex cell boundaries are plotted with a dotted line. The dashed curves correspond to the most common cell boundary descriptions. Note the log scale for the time axis. A consequence of the scaling is that the void area at *t =* 0 is not shown, but can be inferred from Fig. 4

We find that for the OS model, varying the interaction radius significantly impacts the timescale at which the void closes. A repulsion only model results in the quickest closure time for OS models, since cells do not experience any attractive forces. The short-range interaction model closes the void more slowly than the repulsion only model, because of the attractive forces between cells. As cells in the long-range interaction model experience the highest magnitude of attractive force, this model closes the void the most slowly of all of the OS models. These dynamics are shown in Figs. 4(a), (d) and (g) and 5(a). The repulsion only void outlines shown in Fig. 4(a) (OS repulsion) exhibit small tears in the tissue at various times. These tears occur due to localised ‘cell-shoving’. The interaction radius, *r*_max_, dictates the degree of tension within a tissue: for the short and long-range models, increasing *r*_max_ results in higher tension, and hence decreases the rate at which the void closes due to compression. However, even after the void has closed, there are heterogeneities in compression through the tissue, evidenced in Supplementary Video 1, using cell area as a proxy for tissue compression. In the initial stages of void closure there is low compression near the wound edge (this is true across all models and boundary descriptions). Once the void has closed, in the OS models, there is generally a region of lower compression in the centre of the tissue where the void was initially located, and regions of higher compression surrounding this central low compression area. The level and shape of the low compression region and surrounding high compression region depends on the boundary description. In the repulsion case, the low compression region is small and the surrounding high compression region is asymmetrically distributed. In the short-range interaction simulation, the low compression region is slightly elongated in the *y* -direction. Again, the high compression region is asymmetric, with prominent areas in the top-right and left of the low compression region. In the long-range interaction setup, the low compression region is approximately centred with regions of higher compression to the left and right. Heterogeneous stress is also observed in *in vitro* experiments of wound closure, with regions of low stress near the wound edge, surrounded by regions of high stress (Brugués et al., 2014).

For VT models we again obtain distinct behaviour for each of the boundary descriptions. For unbounded tessellations a void cannot be created, since there can be interactions between cells regardless of how far apart they are. This can be seen in Fig. 4(b), showing that if cells are removed, adjacent cells become artificially enlarged to fill the space. Over time, the large cells become smaller and the compressed cells become larger, shown in Supplementary Video 1. In the bounded tessellation case, restructuring of the tessellation causes jump discontinuities in the void area over time, see Fig. 5(b) (VT bounded). For ghost node tessellations, we implement a density dependent ghost node removal, where if a ghost node’s Voronoi area is less than 50% of that at equilibrium, the ghost node is removed from the simulation. As in the bounded tessellation case, ghost node removal leads to discontinuities in void area, as demonstrated in Fig. 5(b). The change in shape of the voids over time for the VT bounded and VT ghost simulations can be seen in Figs. 4(e) and (h), respectively. For both bounded and ghost node tessellations, due to the discrete nature of the model (either from tessellation restructuring or ghost node removal), the void area closure also exhibits a discrete behaviour. We see from the initial dynamics that the void area for both bounded and ghost node tessellations follow similar closure trajectories. However, for the bounded tessellation, once the void is of a certain aspect ratio (where the distance between cell centres on opposite sides of the void is less than 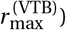, the nature of the tessellation results in the void closing instantaneously. In comparison, the void in the ghost node tessellation case takes longer to close. In each case, once the void has closed the tessellations have a similar form, as seen in Supplementary Video 1. Where the void was originally located, the space has been filled by cells that are larger than those in the surrounding tissue. However, these larger cells are unable to elongate in ways observed *in vitro* because the forces between cells do not take into account orientation or distance from the wound. Surrounding these larger cells is a region of much smaller cells; approximately 0.1 CD^2^ smaller than the cells in the void. Beyond the smaller cells, the cells in the rest of the tissue are of an intermediate size between the larger cells in the centre and the surrounding smaller cells.

The default VM has previously been used to model wound healing due to compression (Nagai and Honda, 2009). The change in shape of the void for the VT simulations is shown in Figs. 4(c), (f) and (i). We see that in both the smoothed boundary model and default model the void closures follow similar trajectories, and close at similar rates. The curved VM requires more time to close the void compared to the other two VMs investigated here, as demonstrated in Fig. 5(c). This is due to the fact that boundary cells have the ability to achieve their idealised area and perimeter more readily due to the extra nodes along the boundary. Hence, once the cells on the boundary have attained their target areas and shapes, the closure dynamics are governed predominantly by tissue compression. In contrast to the OS and VT models, compression is homogeneous throughout the tissue after void closure. In the VM, cells are able to elongate in ways that are not possible in the OS and VT models, as shown in Supplementary Video 1. The explicit incorporation of cell shape in the VM allows the tissue to relax into a configuration where all cells are the same size. Depending on the biological context, this may be desirable behaviour. For example, cell elongation is observed *in vivo* in wound healing experiments and has implications for cell contractility (Brugués et al., 2014). Hence cell-based computational models of wound healing should be able to account for this behaviour.

Overall in this relatively simple scenario of compression driven void closure, the choice of cell boundary description influences void shapes and closure times. In OS models, larger interaction radii can lead to slower void closure rates because cells feel attractive forces towards cells that are further away from the void. An artefact of the VT model is that jump discontinuities occur in the void area over time because of tessellation changes. Furthermore, voids cannot occur in unbounded VT models. In VMs, the greater the degrees of freedom in the cell boundaries at the tissue edge, the more cells are able to deform to reach their target areas and perimeters. For more restrictive cell boundary descriptions, the tissue compensates by enlarging the cells at the tissue boundary more quickly, resulting in faster void closure.

### 3.2 Tissue growth: Choice of model and representation influences tissue density and shape

The second case study we consider is tissue growth. Tissue growth is of considerable interest to both experimental and computational biologists due to the implications for understanding cancer and cellular functions (Kamatar et al., 2020; Mirbagheri et al., 2019). Furthermore, (the simplest) models of tissue growth are amenable to *in vitro* and *in silico* experimentation. We explore how tissues with a free boundary, for example avascular tumours, grow over time due to cell division (Murphy et al., 2022; Galle et al., 2005) (Fig. 2(b)). Such tissues are the subject of many *in vitro* investigations, due to the relative simplicity and low cost of the experimental design (Kamatar et al., 2020; Mirbagheri et al., 2019). These investigations primarily explore the effects of pharmacological intervention on tissue size. In the case where tissue growth is not significantly inhibited, tissues grow to a critical size before cells in the centre of the tissue enter cell cycle arrest (Browning et al., 2021). In the context of tumour growth, the rate of tissue growth is of crucial importance, since fast growing tumours are more likely to be malignant and are associated with lower responses to clinical treatment in non-small cell lung cancer (He et al., 2021). Individual-based models are suitable for studying the effects of individual cell behaviour on the overall tissue size. Here cells in the tissue are able to divide, provided that they are not too compressed, leading to tissue growth. Examples of existing individual-based models of tissue growth are coloured yellow in Fig. 1. We will explore how cell boundary descriptions affect tissue tension in a tissue growing due to cell division. *In vitro* images of avascular tumour growth and a model schematic are shown in Fig. 2(b) and (e).

In our tissue growth simulations, we again begin with a hexagonally-packed tissue, this time centred at the origin, with cell centres located within a radius of five cell diameters of the origin and with free boundaries. These simulations also contain cell division events that cause the tissue to grow, since we do not remove cells from the simulation, causing the tissue to expand into free space. We use a Bernoulli trial incorporating contact inhibition to determine whether a cell will divide. To understand the impacts of stochasticity, we run ten simulations for each model. Parameter values for these simulations are given in Table 3. Sample snapshots of tissues growing over time are shown in Fig. 6. As in the void closure case, tissue boundaries need to be defined for the OS models. We also require tissue boundaries to be defined for the Voronoi tessellation models in this case. Details of the tissue boundary definition are given in Appendix D. The number of cells, circularity of the tissue and proportion of quiescent cells in the tissue over time are presented in Fig. 7. Tissue circularity is calculated as 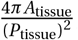, where *A*_tissue_ and *P*_tissue_ are the tissue area and perimeter, respectively. The tissue perimeter is found from the polygon defined by the centres of the boundary cells, and the tissue area is the polygon’s associated area.

**Table 3.**
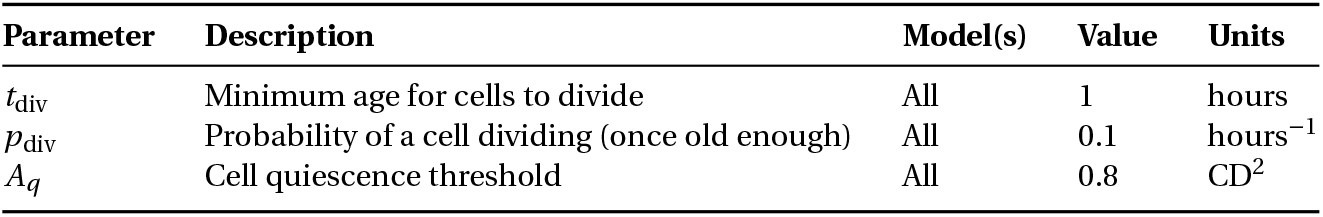
Parameter values used in the tissue growth simulations.

| Parameter | Description | Model(s) | Value | Units |
| --- | --- | --- | --- | --- |
| $t_{\text{div}}$ | Minimum age for cells to divide | All | 1 | hours |
| $p_{\text{div}}$ | Probability of a cell dividing (once old enough) | All | 0.1 | hours<sup>-1</sup> |
| $A_q$ | Cell quiescence threshold | All | 0.8 | CD<sup>2</sup> |

**Table 4.**
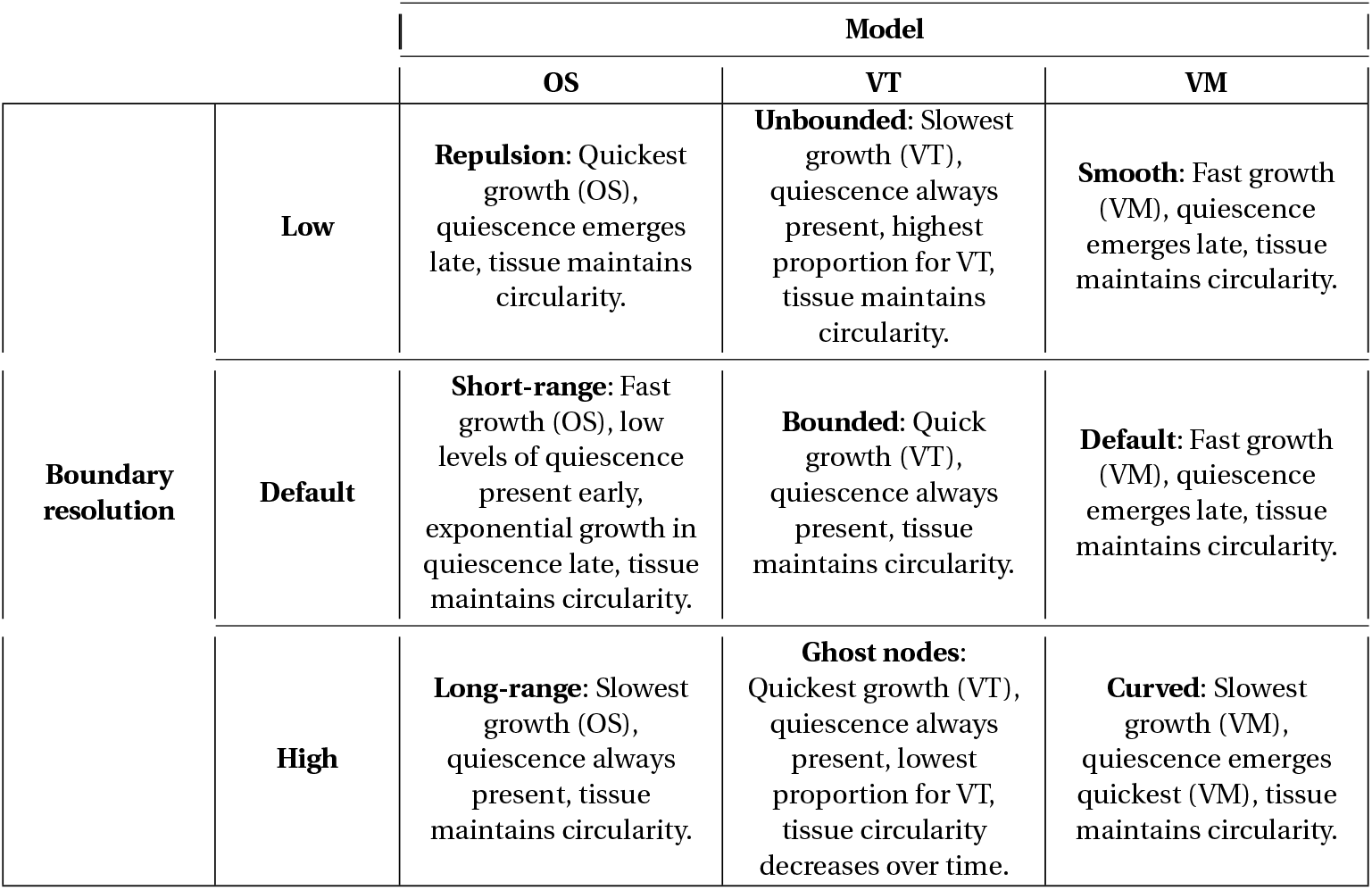
Model comparisons for tissue growth.

|  |  | Model |  |  |
| --- | --- | --- | --- | --- |
|  |  | OS | VT | VM |
| Boundary resolution | Low | **Repulsion:** Quickest growth (OS), quiescence emerges late, tissue maintains circularity. | **Unbounded:** Slowest growth (VT), quiescence always present, highest proportion for VT, tissue maintains circularity. | **Smooth:** Fast growth (VM), quiescence emerges late, tissue maintains circularity. |
|  | Default | **Short-range:** Fast growth (OS), low levels of quiescence present early, exponential growth in quiescence late, tissue maintains circularity. | **Bounded:** Quick growth (VT), quiescence always present, tissue maintains circularity. | **Default:** Fast growth (VM), quiescence emerges late, tissue maintains circularity. |
|  | High | **Long-range:** Slowest growth (OS), quiescence always present, tissue maintains circularity. | **Ghost nodes:** Quickest growth (VT), quiescence always present, lowest proportion for VT, tissue circularity decreases over time. | **Curved:** Slowest growth (VM), quiescence emerges quickest (VM), tissue maintains circularity. |

**Fig. 6.**
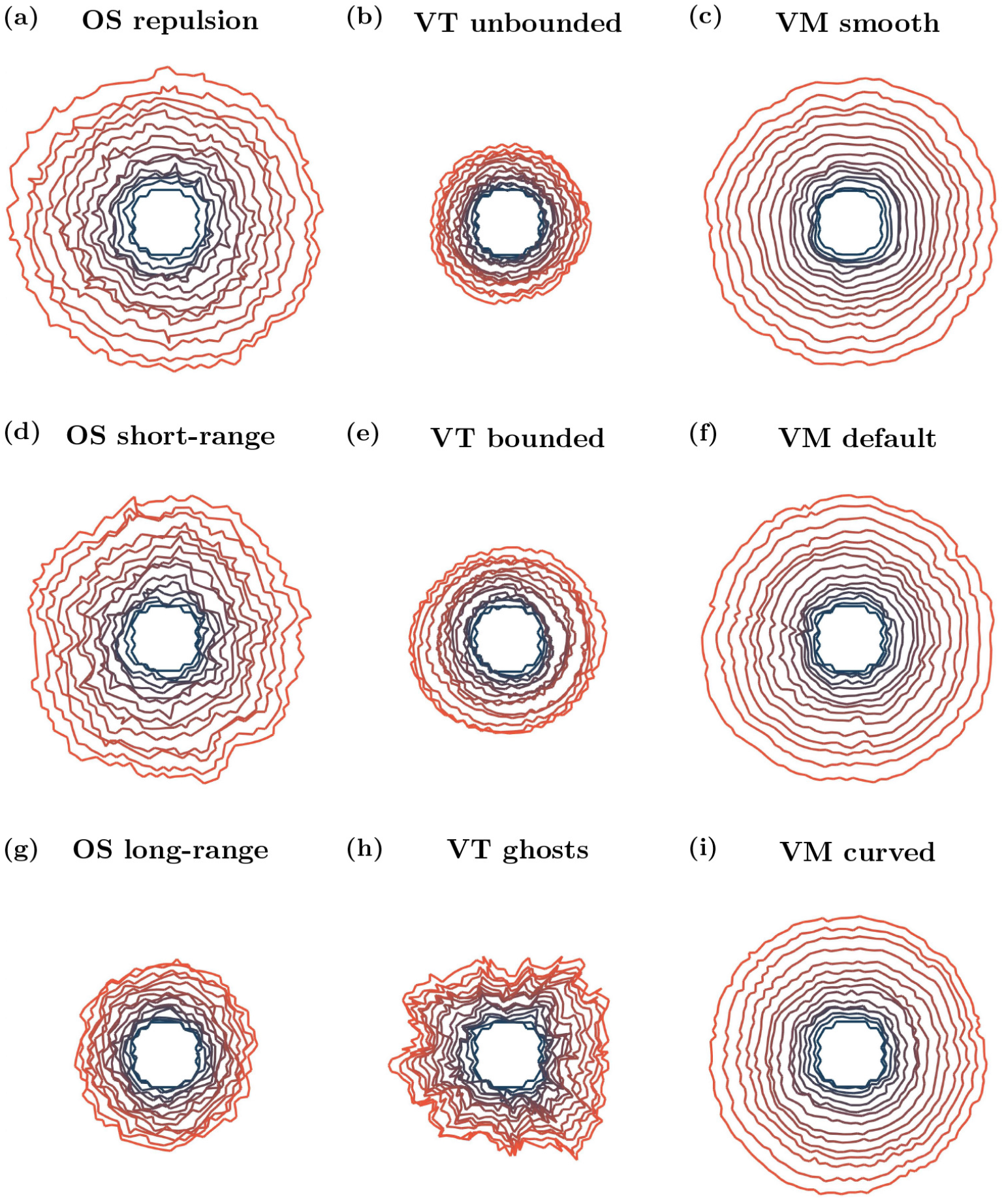
Example growing tissue outlines over time. Initial outline plotted in blue, later times plotted in orange. Outlines plotted every 2.5 hours. Videos are provided in Supplementary Video 2

**Fig. 7.**
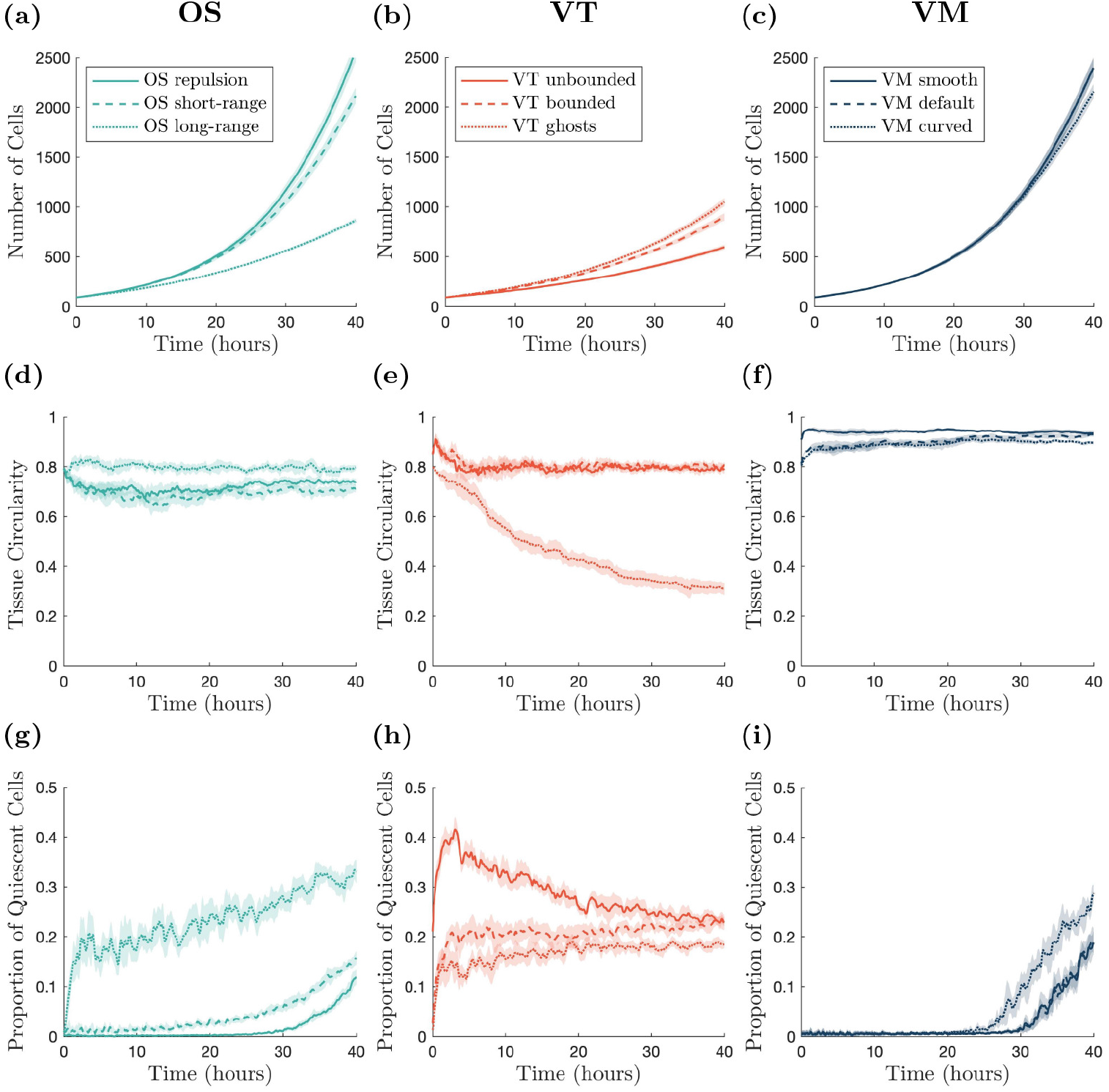
Cell numbers, tissue circularity and cell quiescence over time. (a)-(c) Number of cells within the tissue over time. (d)-(f) Tissue circularity over time. (g)-(i) Proportion of quiescent cells within the tissue over time. Darker lines show averages, shaded regions show 95% confidence intervals for ten simulations. All results are smoothed using a moving average with a sample width of ten data (time) points.

In the OS model, changing the interaction radius affects how large the tissue can grow in 40 hours. For repulsive forces, the tissue is able to grow the largest. Furthermore, over time the tissue becomes most dense near the origin, and less densely packed near the tissue boundary. Increasing the interaction radius decreases the rate of tissue growth, as seen in Figs. 6 and 7(a), (d) and (g), since cells experience greater attractive forces towards each other, and hence do not grow as readily. For long-range interaction forces, the tissue grows most slowly. Fig. 7(d) demonstrates that, in all cases, the circularity of the tissue remains roughly constant over time. In the long-range interaction simulations, quiescent cells appear relatively early in the simulations because of the high tissue density, and the proportion of quiescent cells persists throughout. In contrast, in the repulsion simulations, the proportion of quiescent cells stays close to zero until a quiescent region begins to form at around 26 hours, shown in Fig. 7(g). In the short-range interaction simulations, a small number of quiescent cells develop at early times, but the number of quiescent cells remains low until approximately 15 hours, when the quiescent region begins to develop. Example simulations showing the development of quiescent regions and tissue growth are given in Supplementary Video 2.*In vitro* experiments show that once a tissue is large enough, the cell cycle of the cells in the centre of the tissue arrest, and there is an outer ring of cells whose cell cycle continues to progress (Heinrich et al., 2020). We do not observe this structure in the OS model simulations. However, this may be due to the tissue size not reaching a large enough size.

The type of boundary used in a VT model has a significant impact on the size and shape of a growing tissue with a contact-inhibited cell cycle. Both the bounded and unbounded tessellation simulations are unable to grow as large as the simulations with ghost nodes (Figs. 6 and 7(b), (e) and (h)). However, simulations of tissues with ghost nodes are less circular, with circularity decreasing over time (Fig. 7(e)). To ensure that tissues with ghost nodes remain bounded, there needs to be a sufficient number of ghost nodes surrounding the tissue at the start of the simulation. The number of ghost nodes needed is dependent on the type of tissue growth and how long the simulation runs for. Alternatively, additional ghost nodes need to be added around the tissue as it grows larger. For the ghost node simulations in this work we use eight ghost nodes to the left and right of the tissue, and above and below the tissue (visualised in Supplementary Video 2).In the simulations with ghost nodes, cells are able to interact with fewer neighbours at the boundary, and form finger-like protrusions from the tissue, leading to the tissue becoming less circular. Regardless of the boundary description, quiescent cells appear early in the simulations, as shown in Fig. 7(h). The proportion of quiescent cells is greatest in the unbounded case, although this can be partly attributed to cells around the boundary being quiescent from the start of the simulation, as can be seen in Fig. 7(h). In practice, cells with infinite area at the boundary of the tissue do not have their entire area calculated. Instead their area is calculated using only the finite vertices in the tessellation. The higher proportion of quiescent cells in the unbounded case leads to less tissue growth compared to other simulations. A notable feature of the VT models, which differs from *in vitro* experiments, is that the quiescent cells are relatively uniformly distributed throughout the tissues (except for the boundary effects in the unbounded model). Quiescent cells are ‘isolated’, rather than concentrated in quiescent regions. This is attributed to the linear force used to describe cell interactions, which does not heavily penalise compression, in comparison to the force used in OS. This is shown in Supplementary Video 2, and the feature persists regardless of the strength of the force acting between cells (results not shown).

In the VM, the cell boundary description has a less significant impact on tissue growth and shape compared to the OS and VT models. The smooth and default descriptions exhibit similar rates of tissue growth, as Figs. 6 and7(c), (f) and (i) show, and develop quiescent regions at similar times, as seen in Fig. 7(i)). This is due to the fact that, as the tissue grows, the boundary nodes belonging to a single cell only in the default model make up a smaller proportion of the boundary, and so the default model exhibits the same dynamics as the smooth model. In contrast, the curved VM develops a quiescent region around 5 hours earlier than the smooth and default cases, see Fig. 7(i) and Supplementary Video 2. The earlier emergence of the quiescent region slows tissue growth from around 25 hours for the curved boundary simulations, exhibited in Fig. 7(c). In all VM cases, the region of contact inhibited cells developed approximately in the centre of the tissue as it grew, although the shape of the quiescent regions are asymmetric because of stochastic effects. All boundary descriptions result in highly circular tissues (see Fig. 7(f)). The development of a central region of cells experiencing cell cycle arrest, surrounded by cells progressing through the cell cycle qualitatively agrees with the behaviour exhibited in *in vitro* experiments (Heinrich et al., 2020). However, the emergence of the quiescent region in the VM models occurs when the tissue is small, suggesting that the parameter values in the model need to be calibrated to *in vitro* data.

In the growing tissue example, where cell divisions are dependent on cell size, both cell boundaries and model choice can inhibit tissue growth. Naturally, in the OS model, greater attraction between cells leads to denser tissues, fewer cell divisions and smaller tissue sizes. The VT model is an interesting case, where cell quiescence occurs at early times for all boundary descriptions, resulting in slower tissue growth. In VT models, no central core of dense tissue forms over the time scales that we simulated, rather, isolated quiescent cells appear throughout the tissue. Spatial effects take longer to emerge, if they emerge, in VT models. We have run longer simulations to verify whether the quiescent core structure emerges, but did not observe this behaviour for times up to 60 hours, beyond which point the simulations become extremely computationally expensive because of the large number of cells. Moreover, the level of quiescence observed for the VT models are consistent with the other model descriptions, indicating that a quiescent core does not develop with the same dynamics observed in the other model descriptions. All VM simulations develop quiescent regions. However, the timescale of quiescent region formation varies depending on the boundary description. For smooth and default boundaries, the time scale is approximately the same. For the curved boundary description, the quiescent region forms more quickly.

### 3.3 Tissue collision: Choice of model and representation affects tissue interface shape

Finally, we study the collision of cell fronts that consist of different cell populations. The mechanics of colliding tissues has remained relatively unexplored until recently. Specifically, the dynamics of intersections between tissues with different geometries, cell densities and cell types were only studied *in vitro* and using continuum models in 2022 (Heinrich et al., 2022) (Fig. 2(c)). Given that different tissue types can collide in physiological and pathological scenarios, this may be a fruitful area for future research for biological and computational experimentalists alike. Mechanically, the concept of tissue collisions may involve aspects of void closure and tissue growth previously described. Examples of individual-based models of tissues that grow and collide are shown in blue in Fig. 1. In our work, we allow our colliding tissues to grow via cell division in a confined space, to simulate a much larger tissue. Once a sufficient number of cell divisions have occurred, the tissues will collide. Here we are interested in the shape of the boundary between the two tissues after collision, and how different cell boundary descriptions change this boundary. *In vitro* images of colliding tissues and a model schematic are shown in Fig. 2(c) and (f).

Here, we consider two initially physically separate cell populations growing in a confined region until they collide. Each simulation is started with two columns of cells on the right (labelled cell type A) and left (labelled cell type B) sides of a domain, as shown in Fig. 8. We use a domain which would hexagonally pack a tissue of 14 cells wide and 14 cells high. In this instance, we have reflective boundaries on the left and right and periodic boundaries in the *y* -direction. In the two cell population simulations, we again use a Bernoulli trial in conjunction with contact inhibition to determine whether or not a cell divides. We use the same conditions for division as the growing tissue example. As cells in both populations divide, each tissue grows until the two populations collide. Upon collision, interactions between cells from different cell populations are different compared to interactions with cells of the same type. In the OS model, the elastic interaction constant between cells of the same label remains unchanged, *µ*_*AA*_ *= µ*_*BB*_ *= µ*. However, for cells of a different label, we use a weaker interaction constant, *µ*_*AB*_ *= µ*_*B A*_ *< µ*. Likewise, for the VT model, the spring constant for two cells of the same type remain the same, *κ*_*AA*_ *= κ*_*BB*_ *= κ*, but for cells with different labels it is weaker, *κ*_*AB*_ *= κ*_*B A*_ *< κ*. In the VMs, the adhesion between cells of the same label is given by *γ*_AA_ *= γ*_BB_ *= γ*, while the adhesion between cells of a different label is given by *γ*_AB_ *= γ*_BA_ *> γ*. Parameter values are provided in Table 5, and chosen with the physical interpretation that cells are more likely to interact with other cells of the same label, compared to cells with a different label, to reduce the likelihood of the two cell populations mixing. To determine the location of the interface between the two cell populations, we take the cell centre of all cells which are labelled A, which are in contact with a cell labelled B. We name these cells ‘interface cells’. Interface cells are shown with dark green outlines in Fig. 8. The interface between the two populations is the union of line segments connecting neighbouring cell centres of the interface cells. We note this is not unique, and the equivalent interface would differ slightly if we change the labelling, however, the key results would remain unchanged. An example snapshot of the two tissues after collision at *t =* 40 hours is given in Fig. 8. Since the domain size, and therefore tissue size, is limited in this scenario, we are able to run additional simulations to account for stochasticity. In this case we run 12 simulations per model. Traces of the interfaces for the 12 simulations of each model, and the distribution of the pooled *x*-positions along the interfaces, are given in Fig. 9.

**Table 5.** Parameter values used in colliding tissues simulations.

| Parameter | Description | Model(s) | Value | Units |
| --- | --- | --- | --- | --- |
| $\mu_{AB}$ | Elastic interaction constant between cells with different labels | OS | 5 | CM hours<sup>-2</sup> |
| $\kappa_{AB}$ | Spring constant between cells with different labels | VT | 5 | CM hours<sup>-2</sup> |
| $\gamma_{AB}$ | Adhesion between cells in different tissues | VM | 2 | CM CD hours<sup>-2</sup> |

**Table 6.**
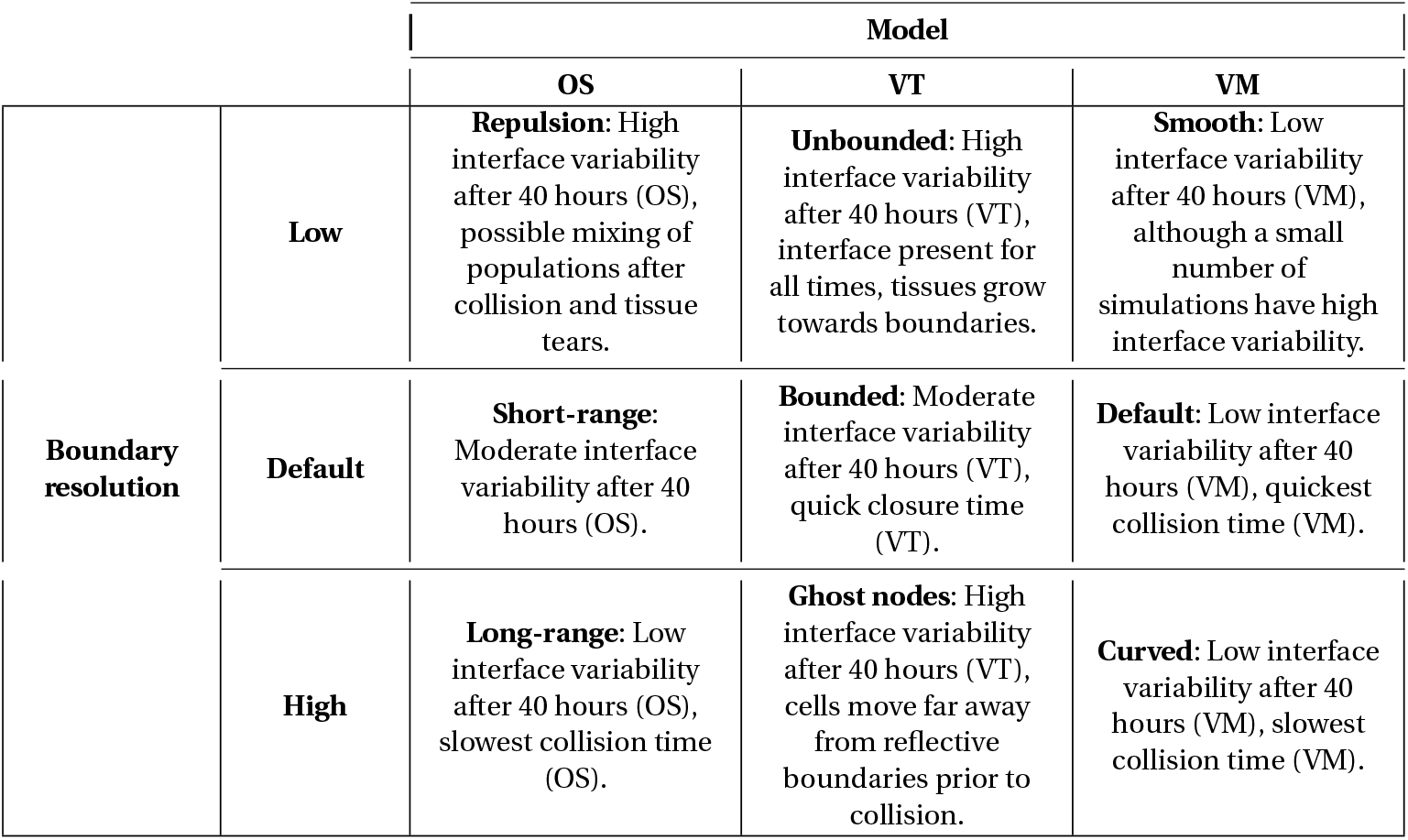
Model comparisons for tissue collisions.

**Fig. 8.**
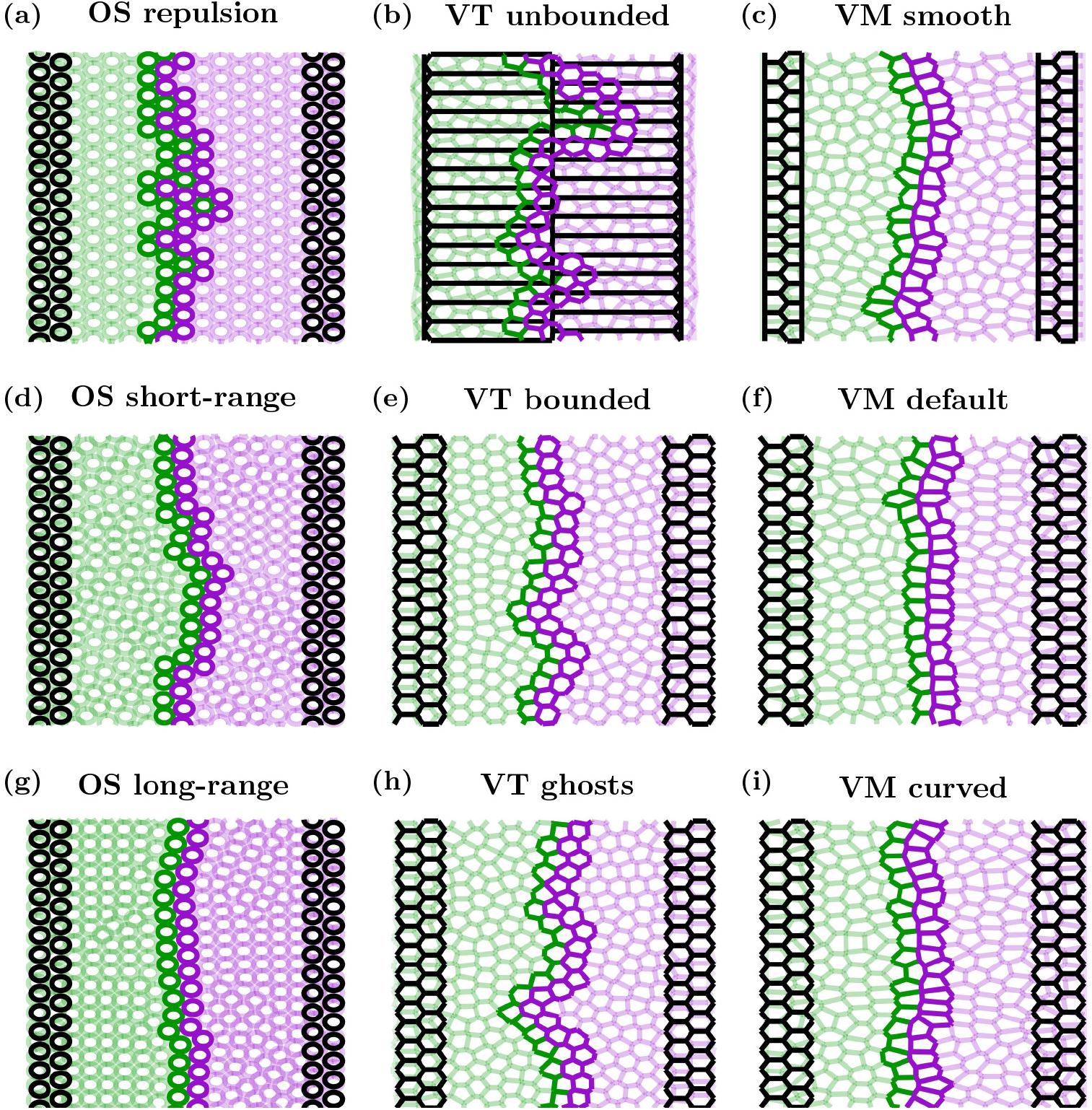
Cell outlines for example tissue collision simulations. Initial conditions are shown in black, final frames (at *t =* 40 hours) are shown by a green population on the left (cell label B) and purple population on the right (cell label A). Cells at the interface of the two populations are indicated via darker outlines. For clarity, the cells in the OS model have been plotted with a size of 0.95 CD. Videos of each simulation are provided in Supplementary Video 3

**Fig. 9.**
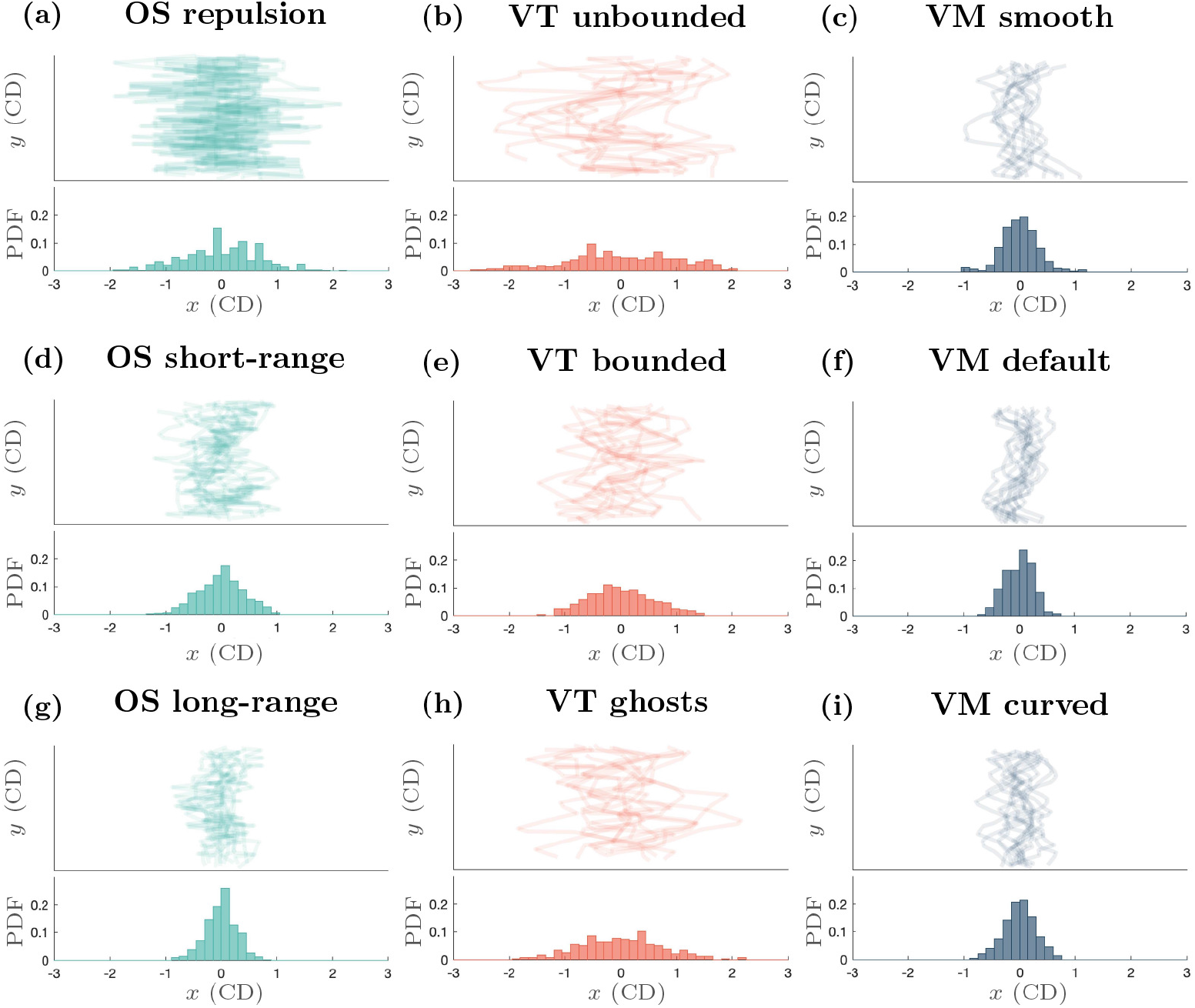
Tissue collision interface structure. Top panels: interface plots (*x*-*y* axes) for the 12 individual simulations of each model at *t =* 40 hours, after the two populations have collided. Bottom panels: histograms showing the probability distributions of the pooled *x*-positions of the interface between the two populations for the 12 simulations

In the OS models, we find that larger interaction radii lead to straighter interfaces between the two cell populations once they collide. As mentioned in the previous examples, larger interaction radii result in denser tissues, which in this case gives rise to a less jagged interface, since cells experience more attraction to their source tissue. This can be seen in Figs. 8 and 9(a), (d) and (g). In Fig. 9(a), (d) and (g), we see lines with greater variation in the *x*-position for the repulsion case, and much smaller variation in the long-range interaction case. This is also reflected in the histograms showing the distributions of the *x*-position along the interface, with the spread in the distributions of the interface decreasing as we increase *r*_max_. In the repulsion case, there are instances where cells become detached from their source tissue, as can be seen by the isolated (dark) green cell, surrounded by (dark) purple cells approximately halfway up the tissue in Fig. 8(a)) and Supplementary Video 3. Changes in the shape of the boundary of each (pre-collision, isolated) tissue occur due to the stochastic cell division events, and these stochastic changes in tissue boundary are amplified in the repulsion case compared to the other two cases, since mature cells experience no attractive forces towards each other. However, once the tissues collide, the variation in the *x*-positions along the interface decrease as the tissue relaxes. As in the void closure case, the repulsion-only tissues collide first, followed by the short-range interaction tissues, with the long-range interaction tissues colliding last, shown in Supplementary Video 3.

For the VT models, the unbounded tessellations have the highest variation of *x*-position at the interface. The bounded tessellations had the least variation in *x*-position along the interface, as evidenced in Figs. 8 and 9(b), (e) and (h). We note that there is always an interface between the two cell populations in the unbounded tessellation and the tissues grow outwards, rather than growing towards each other (see Supplementary Video 3), however, we only consider the shape of the interface between the populations at *t =* 40 hours. In the bounded and ghost node simulations, the tissues also move away from the reflective boundaries throughout the simulation, shown in Supplementary Video 3. This is most notable in the ghost node case, where the tissues form highly irregular shapes until the domain is filled. Similarly to the void closure scenario, but in contrast to the tissue growth scenario, the ghost node simulations took the longest to fill the domain in the tissue collision simulations.

In the VMs, the default and curved boundary cases have similar spread in *x*-position along the interface, as shown in Figs. 8 and 9(c), (f) and (i). In the smooth boundary case there is some additional variation in *x*-position of the interface, as seen in Figs. 8 and 9(c). However, the shape of the tissue boundaries before collisions are broadly consistent across cell boundary descriptions. The slight increase in frequency of *x*-positions near *±*1 CD in the smooth boundary case seen in Fig. 9(c) can be explained by the regions of purple (or green) tissue extending into the green (or purple) tissue and forming a flat boundary near *x = ±*1 CD in some simulations. This can be seen in the interface tracings in the top panel of Fig. 9(c). Similarly to the previous two biological scenarios considered, the choice of cell boundary on the tissue edge affects the time scale of the tissues colliding, with smooth simulations again being the fastest, but on a similar time scale to the default boundaries, and the curved boundaries being the slowest, as demonstrated in Supplementary Video 3.

For the tissue collision example, we find the shape of the interface between two cell populations varies depending on cell boundary description. For OS models, we see that long-range interactions result in less variable interfaces, whereas repulsion-only models have greater variation. As in the void example, unbounded VT models are inappropriate for modelling collisions between tissues, as cells interact with other cells that are too far apart, which would be physically unrealistic. Of the remaining two boundary descriptions for the VT model, the ghost node description has a more variable interface than the bounded tessellation description. As in the previous biological examples, we observe that the shape of the interface is least sensitive to the boundary description in VMs, however, the time scale of the tissues colliding can vary depending on the boundary description (not shown).

## 4 Discussion

When simulating biological tissues with individual-based models, there are many considerations to take into account, such as the complexity of the individual-based model and the biological relevance of the model. Off-lattice models are explicitly informed by physical principles that determine individual cell behaviour and interactions between cells. As such, these models can readily investigate mechanical hypotheses about cell behaviour. Recent work into off-lattice individual-based models have proposed modifications to cell boundaries, particularly at tissue boundaries, that increase the complexity of the model (Ishimoto and Morishita, 2014; Kachalo et al., 2015; Mosaffa et al., 2020; Germano et al., 2022). In this work, we use three common individual-based models; the OS, VT and VMs - and three cell boundary descriptions for each model to investigate the impact of cell boundary descriptions on tissue behaviour in three biological scenarios. In total 27 simulation setups were explored, with multiple simulations run for setups with stochastic effects.

The OS model is known to result in (sometimes) physically unrealistic artefacts because of the way cell interactions are defined (Pathmanathan et al., 2009; Osborne et al., 2017). Specifically, if the interaction radius is large enough, cells may physically interact with cells that they are not in direct contact with. This can result in tissues being artificially dense, as seen in our long-range interaction simulations. If cells are able to emit chemical cues to other cells, as opposed to mechanical cues, that can pass by intermediate cells without distortion, then long-range OS models may be appropriate to model such interactions. One approach to resolve this artefact, is to use a Delaunay triangulation to define cell neighbours (as in the VT case), but with the OS force function, Equation (3) (Mathias et al., 2020). An advantage of the OS model is that it is the least computationally expensive model of those considered here. This makes the OS model more amenable to simulating larger tissues, provided the assumptions of the model hold. The OS model is useful for modelling tissue-scale behaviour, despite no explicit representation of cell shape. However, as cells are represented by points in space, it is less useful for analysing specific tissue boundary shapes, as cell boundaries are not explicitly described by the model. Detailed descriptions of how we describe tissue boundaries in the OS model for the void closure and growing tissue cases are given in Appendices C and D, respectively. However, alternative choices could be made that result in different simulation snapshots and area curves in the internal void and tissue growth examples. It is clear that the interaction radius of the OS model has a significant impact on simulation results across all three biological scenarios that we studied. Generally, the most commonly-used interaction radius, the short-range interaction case, is a reasonable choice. In the void example, the void with short-range interactions is able to close in a reasonable time without causing tissue tears (Figs. 4(d) and 5(a)). However, in the tissue growth example, the repulsion model is better able to capture spatial effects on cell division behaviour. Specifically, while the tissue is small there is negligible compression on cells, but once a critical mass is reached, the number of compressed (quiescent) cells begins to increase exponentially (Fig. 7(g)). In contrast, in the tissue collision example, we see that the long-range interaction model better describes *in vitro* experiments, with less variation at the tissue interface, seen in Fig. 9(g).

VT models are more computationally expensive than their OS counterparts. However, cells cannot physically interact ‘through’ other cells, and therefore VT models more realistically represent mechanical interactions between cell neighbours. In all biological scenarios we explore, VT models exhibit unique, and sometimes surprising, behaviour. In the void closure scenario, voids either cannot be defined, as in the unbounded VT case, or tessellation changes, such as ghost node removal, cause unrealistic deformations in void shape. In the tissue growth example, the presence of quiescent cells from early times in the simulations in all cases prevent the tissue from growing as quickly as in the other models, regardless of the strength of the interactions between cells. This suggests that the VT model may not be appropriate for capturing cell compression in growing tissues, unless coupled with chemical or other signalling pathways. Unbounded VT models are often unphysical, as shown in Figs. 4, 6 and 8(b), and generally should be avoided. Bounded VT models had the least variation in tissue shape in all biological scenarios, compared to the other VT cell boundary descriptions. This is exemplified in Figs. 7 and 9(e). Tissue shape is highly irregular when ghost nodes are used, and indeed in the tissue growth examples, ghost nodes can ‘infiltrate’ the tissue. These artefacts can be removed by deleting ghost nodes that are not connected to any other ghost nodes in the Delaunay triangulation, however, sometimes ghost nodes penetrate the tissue in pairs or triples (and possibly larger configurations). Handling the artefacts arising from these cases requires checking whether a group of ghost nodes is surrounded by real cells and adds to the computational cost of the system. An additional subtlety to consider when using ghost nodes is the density of the ghost nodes. In our simulations, we assume ghost nodes have the same density as real cells. If ghost nodes are more or less dense than the real cells, then the shape of the tissue will be altered and overall dynamics of the system may be impacted. For example, in the void closure scenario, if the ghost node density is greater than the density of the real cells, the discontinuities in the void area over time curves may not be so pronounced, provided that ghost node removal was defined appropriately, and not all of the ghost nodes are removed simultaneously. The criteria for removing ghost nodes can similarly impact the void area over time curve. In the tissue growth example, if the ghost nodes are more dense than the real nodes then there may be more infiltration of ghost nodes into the tissue. The effect of the density of ghost nodes is less significant in the tissue collision scenario, since we are interested in how the tissues behave after they collide, when there are no ghost nodes between the two tissues. Still, ghost node density will affect both the time at which the tissues collide and the shape of the interface between the two tissues until the overall tissue reaches confluence. There are several choices that must be made when using ghost nodes which can result in unexpected computational artefacts if not appropriately made.

The VM does not suffer from the same tissue shape artefacts as the OS or VT models, as the tissue boundary is explicitly defined in the VM. However, the VM is the most computationally expensive model considered here. A portion of the computational cost can be attributed to the forces acting on polygonal vertices, rather than cell centres. This results in the number of force calculations performed in a VM being approximately six times the number performed in the OS and VT models. This is magnified further for the curved boundary model where additional nodes are introduced. The greater cost, however, arises from the manual cell rearrangement and mesh restructuring operations required in the VM. Additionally, the VM is highly parameterised, and great care must be taken when selecting parameters, as the resulting dynamics are highly sensitive to model parameters. In the biological scenarios we consider here, the intersection of tissues often requires additional mesh restructuring operations, which are detailed in Appendix B. The more restructuring operations that need to be checked and performed, the more expensive simulations become. The VM is the least sensitive to changes in cell boundary description, with the smooth and default descriptions behaving similarly across all biological scenarios. The curved model is the most computationally expensive of all the models here. Furthermore, tissue behaviour evolves on a longer timescale in the curved model compared to the smooth and default cases in the void closure and tissue collision examples. The additional degrees of freedom in the cells on the boundary in the curved model mean that the boundary cells do not have to rely on interactions with their neighbours to deform and attain their target areas and perimeters. Arguably, this makes the curved VM the most ‘realistic’ model we have considered in this work.

Another confounding factor to consider during model selection is the typical run-time of the simulation. In Table 7 we provide typical run-times required for each simulation presented here. Note that the run-time includes all calculations required to obtain the results presented in the Results section (for example, void area calculations or tissue circularity calculations that introduce computational overheads beyond model implementation). We note these results should be compared within models, rather than across models (i.e. different variants of the OS model can be compared, but the OS model should not be compared to the VT or VM), as model implementation significantly contributes to run-time.

**Table 7.** Typical run-time. As measured in seconds, on single core of a 2.10GHz Intel Xeon E5-2683 v4 CPU, with 240GB of RAM. Final cell numbers are given in parentheses for the tissue growth and tissue collision scenarios. Cell numbers are constant for the void closure simulations. For the OS model, the low boundary resolution is the repulsion model, the default boundary resolution in the short-range interaction model and the high boundary resolution is the long-range interaction model. For the VT model, the low boundary resolution is the unbounded model, the default boundary resolution is the bounded model and the high boundary resolution is the ghost node model. For the VM, the low boundary resolution is the smooth model, the default boundary resolution is the default model and the high boundary resolution is the curved model.

|  |  |  |  | Model |  |  |
| --- | --- | --- | --- | --- | --- | --- |
|  |  |  |  | OS | VT | VM |
| Biological scenario | Void closure<br>(204 Cells) |  | Low | 3,700s | 510s | 1,800s |
|  |  |  | Default | 3,800s | 570s | 2,000s |
|  |  |  | High | 3,700s | 570s | 2,200s |
|  | Tissue growth | Boundary resolution | Low | 30,500s<br>(2,570 cells) | 6,400s<br>(600 cells) | 1,200s<br>(2,400 cells) |
|  |  |  | Default | 22,400s<br>(2,120 cells) | 1,400s<br>(900 cells) | 1,400s<br>(2,400 cells) |
|  |  |  | High | 5,100s<br>(860 cells) | 6,500s<br>(1,050 cells) | 116,700s<br>(2,150 cells) |
|  | Tissue collision |  | Low | 210s<br>(283 cells) | 2,900s<br>(285 cells) | 530s<br>(251 cells) |
|  |  |  | Default | 240s<br>(302 cells) | 820s<br>(298 cells) | 550s<br>(256 cells) |
|  |  |  | High | 280s<br>(330 cells) | 1,030s<br>(296 cells) | 21,880s<br>(256 cells) |

For the void closure scenario, we observe that the run-time does not vary significantly with an increasing boundary resolution. This is due to the fact that the number of cells remains constant within this scenario, and computational complexity is roughly consistent.

For the tissue growth scenario, using the OS model, we observe that the run-time decreases with an increase in boundary resolution. This is explained by the final number of cells within (Fig 7, which shows the final number of cells decreasing with increasing boundary resolution. For the VT model, we find that both the ghost node and unbounded models perform comparably. In contrast, due to an increased computational load in calculating cell bounds, the bounded model requires a significantly longer run-time, despite the higher number of cells in the ghost node model. We see that the curved VM model has a run-time that is two orders of magnitude larger than the other VM models, even though the number of cells is similar. This difference is better understood by noting that the run-time for VM depends on the number of nodes (vertices), rather than the number of cells, and the number of nodes (vertices) in the VM curved model grows rapidly with the growing tissue.

Lastly, for the tissue collision scenario, we see that the OS models all have comparable run-times, with the low resolution models having the shortest run-time, and run-time increasing with boundary resolution. This is because the increased boundary resolution also increases tissue compression, delaying the tissue reaching confluence. For the VT models, we see that the run-time also increases with boundary resolution. The increase in run-time of the VT model is due to the computational complexity of performing the tissue bounding (for the VT bounded) and removing ghost nodes (for VT ghosts). Lastly, the VM run-time also increases with boundary resolution, with a significant increase of two orders of magnitude for the VM curved model. However, even though the tissue boundary is growing, it is not growing analogously to the tissue growth case. Instead, the significant increase in run-time is due to node-node and node-edge collisions as the two tissues collide, requiring resolutions and introducing additional computational complexity.

While we have considered boundary effects in this work, an avenue of individual-based models that is yet to be fully explored is that of model stability, parameter values and quantitative comparisons with *in vitro* or *in vivo* data. Given the interactions between multiple cells and the computational costs of these models (even the simplest ones), these issues are not straightforward to resolve and have been overshadowed by the drive to ensure that the models qualitatively describe biological data. Future investigations into individual-based models should investigate which parameter values ensure the mathematical stability of these models. Moreover, understanding the range of parameter values that result in biologically relevant simulations and quantitative analyses of how closely simulations compare with biological data is highly desired, as parameter fitting for these models is a computationally expensive task. Efforts have been made to resolve parameter values for both on-lattice and off-lattice models (Jagiella et al., 2017; Kursawe et al., 2018). As technologies advance, the opportunities for parameter exploration are expanding through the use of high-performance computing, parallelisation, and access to increasing volumes of high-quality data (Montagud et al., 2021). Lastly, an approach which would significantly advance the field of individual-based models is a natural way to move between the cell-scale, tissue-scale and organ-scale. Ideally, this would be in the form of a multi-scale modelling framework.

We have restricted our exploration in this paper to two-dimensional examples. An investigation into three-dimensional tissue boundaries would further elucidate the influence of cell boundary descriptions on computational studies of biological phenomena. We expect that the two-dimensional artefacts, such as differing time scales for biological behaviour and observations of tissue shape effects will be exaggerated in three dimensions. Of course, moving into three dimensions adds to the computational complexity. The OS and VT models are both relatively straightforward to extend to three dimensions. However, the VM involves careful consideration of cell rearrangements and the intersections of tissues, which is already quite complex in two dimensions. As such, we leave this for future work.

## 5 Conclusion

In this paper we have demonstrated that the choice of cell boundary descriptions in individual-based models can significantly influence behaviour at a tissue scale. In our case study of a void within a compressed tissue, we observe that the cell boundary description changes the time scale of void closure. In each case, we find that the ‘simplest’ cell boundary description (repulsion-only for the OS model, unbounded tessellations in the VT model and smooth boundaries in the VM) leads to the fastest void closure. In the case of the unbounded VT model, a void cannot even be (naturally) defined. In contrast, the most complex cell boundary descriptions (long-range interactions for OS models, ghost nodes for the VT model and curved boundaries in the VM) have the longest void closure times. However, in the case of the VM, this may be the most realistic representation of void closure, since cells on the tissue boundary are able to deform more. In our case study of tissue growth, we find that all VT models have relatively uniform cell compression throughout the tissue. This is similarly true of the long-range interaction OS model. However, this uniformity of compression is not observed in *in vitro* experiments of growing tissues. In comparison, all VMs, and the repulsion and short-range OS models were successfully able to characterise cell compression as the tissue grew. In our case study of tissue collision, we find that the cell boundary description influences the shape of the interface between the two populations after they have collided. The VT models show the greatest variability of the *x*-coordinates of the interface between the two populations and the VMs showed the least variation. The variation in the OS model depends strongly on which cell boundary description is used.

Overall, the OS model is the most sensitive to changes in cell boundary description, since this has a direct effect on the forces acting between cells. The VM was the least sensitive to changes in cell boundary description, especially with regard to tissue shape in the tissue growth and tissue collision examples. However, the curved VM evolves on a different time scale to both the smooth and default VMs in all biological scenarios investigated. Finally, the VT model exhibits different artefacts depending on the cell boundary description used. Using the unbounded description, voids cannot be described, and space between tissues cannot exist. In the void example, the nature of tessellations results in jump discontinuities in the void area over time. In the tissue growth example, tissues with ghost nodes exhibit highly irregular tissue shapes. The VT model is useful because of its ability to appropriately define cell shapes in confluent tissues and define neighbours to calculate forces between cells. However, this work is a cautionary tale of the artefacts that can arise when using VT models, or indeed OS or VMs; these issues must be carefully considered when interpreting simulation results.

Individual-based modelling is relatively new in the context of biological modelling and there are many avenues for exploration. Future work is necessary to more carefully analyse the stability of individual-based models and the parameter space where these models can provide relevant biological insight. These future investigations will ideally be conducted in three dimensions, with the caveat that the VM becomes much more complex in the three-dimensional case and is currently computationally infeasible for such a study.

## Supplementary information

Supplementary Video 1: Void closure https://drive.google.com/file/d/1jAZ-qxptpvPp91WX3qapL2xakvJhGNvl/view?usp=sharing.

Supplementary Video 2: Growing monolayer https://drive.google.com/file/d/1DUUHEb4BPUf6oTdDJ-RPhEAVUxE65F3Z/view?usp=sharing. Brown cells are able to divide, red cells are quiescent.

Supplementary Video 3: Competing populations https://drive.google.com/file/d/1t0XvJ5vO4dm6Rfp9mweMrhjPPiiZYvl-/view?usp=sharing.

## Acknowledgments

This research was supported by Australian Government Research Training Program (RTP) Scholarships, awarded individually to D.P.J. Germano and A. Zanca. S.T. Johnston’s research is supported by the Australian Research Council (DE200100998). J.A. Flegg’s research is supported by the Australian Research Council (DP200100747, FT210100034).

## Declarations

Availability of data and materials: All data generated or analysed during this study are included in this published article and its supplementary information files

## Appendix A Bounded VT model

In order to implement bounding of cells in the VT model we extend the Delaunay mesh by adding a new node perpendicular to any external edge (i.e inside wound void or outside growing domain. 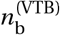 evenly spaced nodes are added a distance 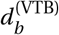 from the boundary edge, See Fig. A1. Note that if one node is added it is placed in the centre of the edge otherwise two nodes are placed at the end and the remaining nodes are spread evenly. Here we use either one (void and competing populations) or two nodes per edge (monolayer). If any new node being added is within a given threshold of any existing nodes (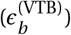_)_) then it is not added this is to avoid a degenerate case of multiple nodes occupying the same location. The Voronoi regions of all cells are then calculated which are now all bounded. Once calculated all the extra image nodes are removed and we continue on as for the non bounded case. An example of the image nodes and resulting bounded Voronoi regions is given in Fig. A1. The parameters used in the model are provided in Table A1.

**Fig. A1.**
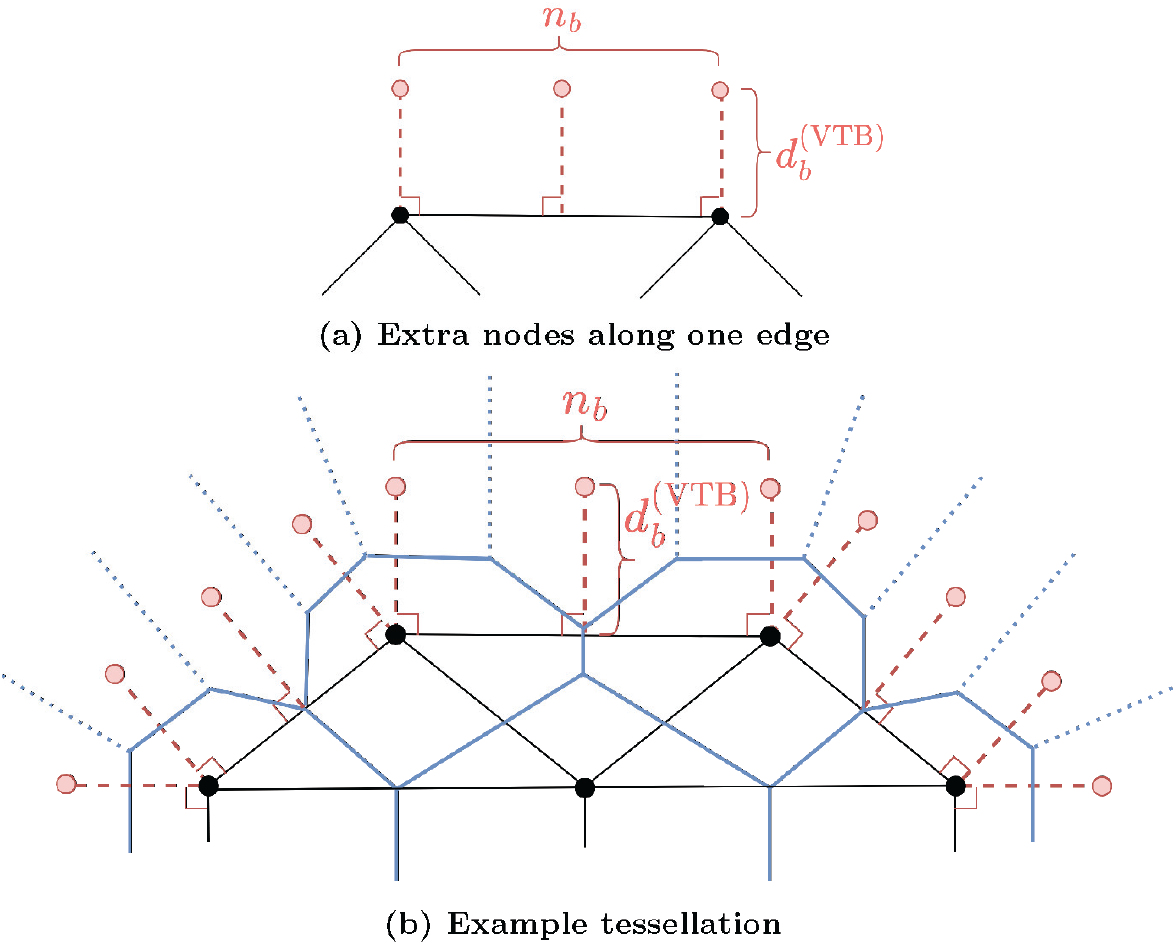
How to allocate extra nodes in the bounded VT simulations

**Table A1.** Parameter values used in the bounded VT model

| Parameter | Description | Value | Units |
| --- | --- | --- | --- |
| $n_b^{(VTB)}$ | Number of bounding nodes | 2 | – |
| $d_b^{(VTB)}$ | Distance between existing edge and added nodes | 1 | CD |
| $\epsilon_b^{(VTB)}$ | Minimum distance between existing and added nodes | $10^{-5}$ | CD |

## Appendix B Vertex model boundary node remodelling

One of the difficulties around using a vertex model is having to resolve various node-node or node-edge collisions. Here we briefly outline the key remodelling decisions implemented to complete this work.

### B.1 Boundary node-boundary node collision with a neighbouring node in common

An instance which causes issues if gone unchecked is when two boundary nodes who share a common neighbouring node collide, as shown in Fig. B2. If the boundary nodes become too close (depending on the cellular boundary description of vertex model), then the nodes are merged with the neighbouring node in common, which is then placed at the midpoint of the two original nodes.

**Fig. B2.**
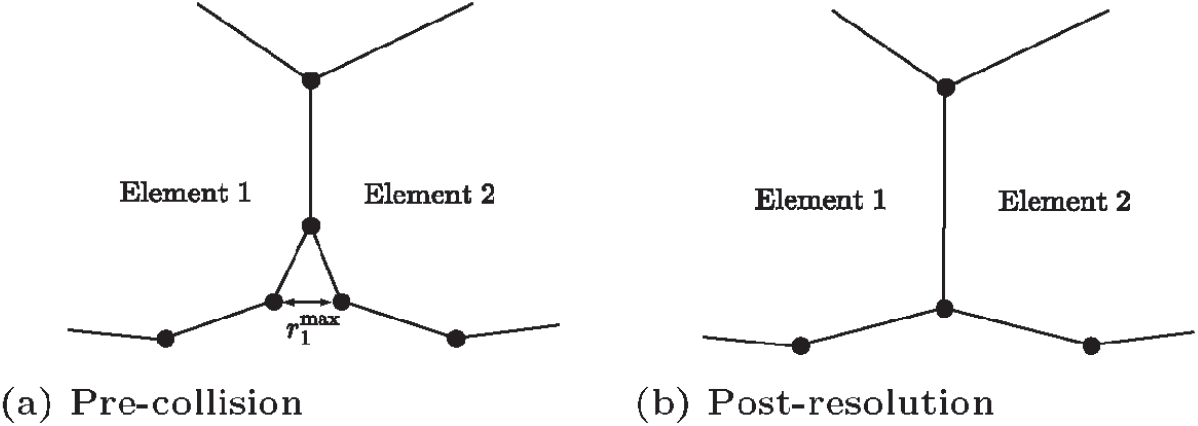
Resolution of boundary node-boundary node collision with a neighbouring node in common. The neighbouring node in common is shared by elements 1 and 2, with the nodes are colliding

### B.2 Boundary node-boundary edge collision

As we have introduced a curved vertex model, we can have boundary edges being very small, in comparison to a traditional vertex model. Therefore, in a previous work, such as (Fletcher et al., 2013), when a boundary node intersects a boundary edge, the nodes making the edge would be moved apart, to make room for the incoming boundary node. However, as our edges here are too small, this causes issues, such as collisions with other nodes, or collisions into other elements. To prevent this occurring, if a boundary node becomes too close to a boundary edge, we merge the boundary node into the element that edge belongs to. This joins the two elements together. The boundary node is placed at the midpoint of its original position and the closest point of the edge. A schematic is given in Fig. B3.

**Fig. B3.**
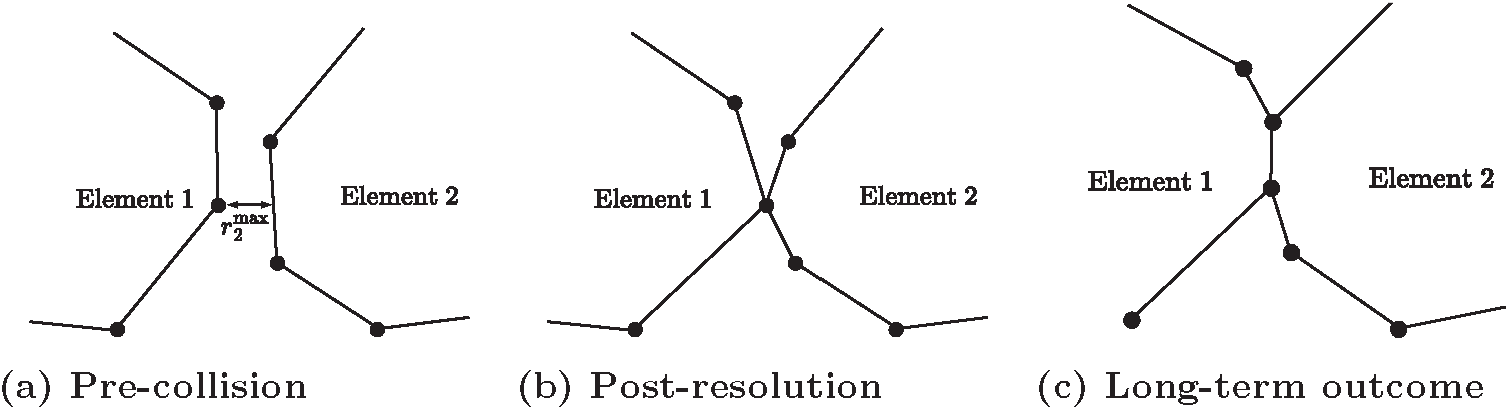
Resolution of boundary node-boundary edge collision. Here, the node that is too close to the edge is added to the element belonging to the edge, joining the two elements together

### B.3 Preemptive boundary T1

A conventional T1 swap in a vertex model is performed when the edge between two interacting elements becomes too small, and the edge of interaction is switched to the adjacent neighbouring elements, as described in (Fletcher et al., 2013). Here, we do not have the exact scenario as above as the nodes interacting are boundary nodes. In this instance, when two boundary nodes which belong to two separate elements are sufficiently close to each other, we perturb the nodes perpendicular to their pairwise displacement, thus preventing the nodes from colliding and possibly merging, creating a rosette. A schematic is given in Fig. B4.

**Fig. B4.**
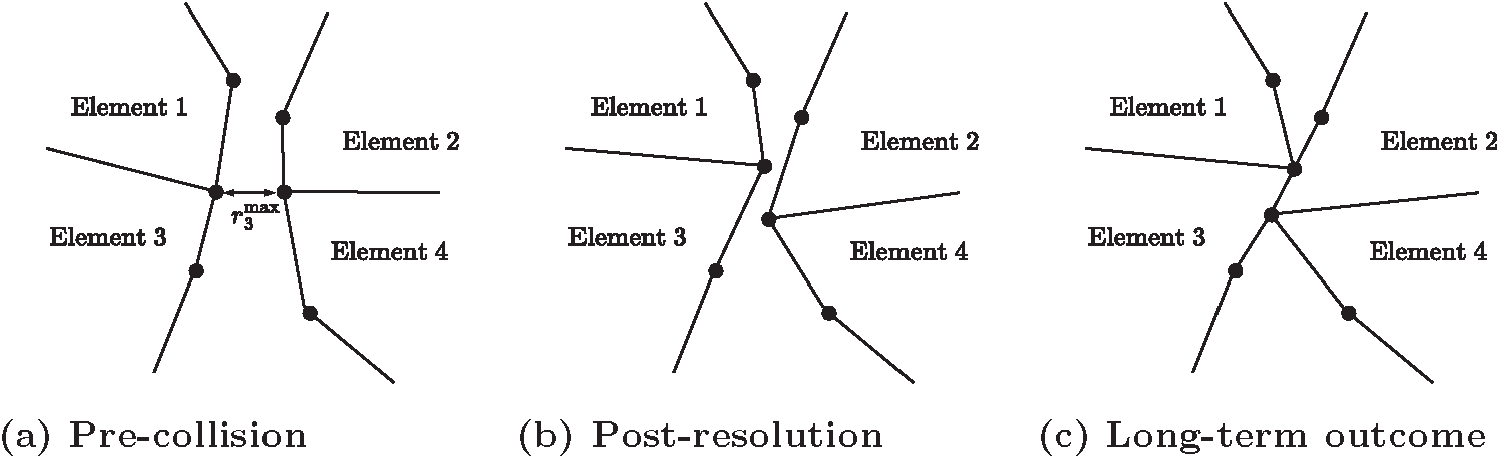
Resolution of boundary node-boundary edge collision. Here, the node that is too close to the edge is added to the element belonging to the edge, joining the two elements together

### B.4 T2 void removal consisting of three boundary nodes

We often have the case of three elements closing a gap to all meet each other. When this happens, an internal void consisting of 3 boundary nodes forms. In previous work, a void of this nature would be removed via a series of T3 swaps, as the tissue is undergoing large amounts of compression. However, in our scenarios, we rarely have this high degree of compression, and therefore the series of T3 swaps do not occur. To compensate for this, when the void area, *A*_*V*_ is sufficiently small, 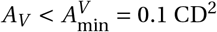, we remove the void by merging the 3 boundary nodes at their centre of mass. A schematic of this scenario is given in Fig. B5.

**Fig. B5.**
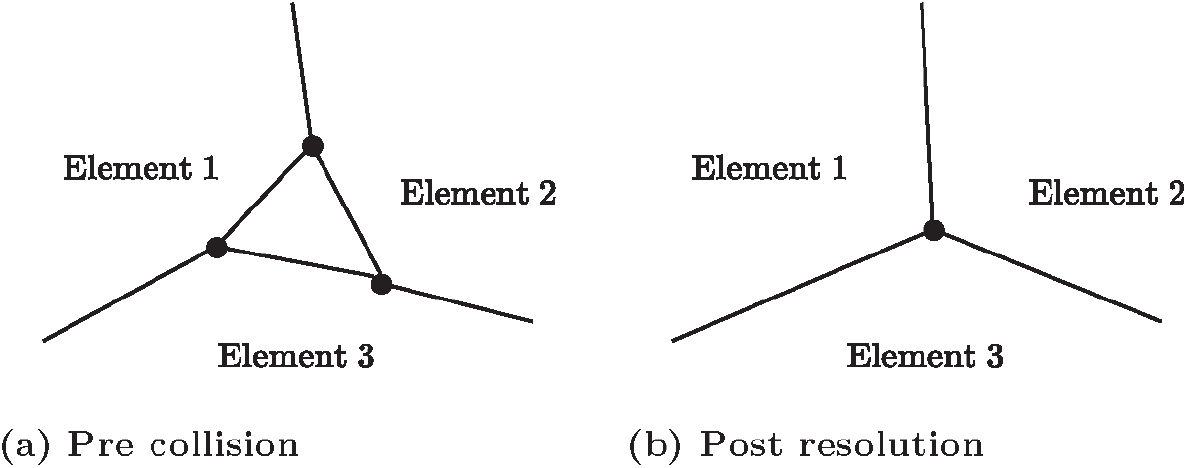
Resolution of void consisting of boundary nodes. When the void is sufficiently small, we remove the void by merging the 3 boundary nodes at their centre of mass. The merged node is now an internal node

### B.5 Boundary node labelling

As the fundamental change we are implementing is how the boundary is modelled, we needed a consistent method for tracking either the boundary or internal status of a node. Previously, vertex models do not contain highly dynamic boundaries, and as such, the status of a node has not been important. To rectify this, each time step, we update the status of each node, labelling it as either boundary or internal. This is done by checking how many elements a node belongs to, and compare that to how many node neighbours a node has. If the node has more neighbouring nodes than the number of elements it belongs to, it is a boundary node.

## Appendix C Calculating void area

Finite Voronoi tessellations and vertex models naturally account for cell boundaries. For these models, the area of the void is the difference in the area of the tissue and the total area the cells occupy. By definition, in an infinite Voronoi tessellation, removing cells does not create a void, so we can never model a void. Overlapping spheres models do not have explicitly defined boundaries with which to calculate void areas. Therefore, a method for determining the void area for overlapping spheres is required. To do this, we first layout a discrete pixel background mesh, of resolution sufficiently small to obtain the desired accuracy. We then assign a cell-pixel interaction radius, *r*_cell-pixel_ *=* 0.50 CD, to each cell. If a pixel falls within a cell’s cell-pixel interaction radius, it does not contribute to the void area, otherwise, it does. A depiction of the calculation is shown for the overlapping spheres model in Fig. C6.

**Fig. C6.**
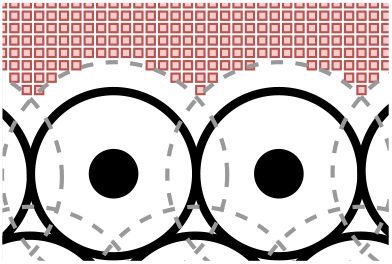
A depiction of the void area calculation for overlapping spheres. The solid black dot represents the cell’s centre, the solid black circle the cell’s boundary and the dashed grey circle the cell-pixel interaction radius, *r*_cell-pixel_. The red squares are the discrete pixels contributing to the void area

## Appendix D Determining growing monolayer boundary

Similarly to the void area problem, seeing as overlapping spheres models, unbounded Voronoi tessellations and finite Voronoi tessellations have no defined boundary, we here introduce a method to calculate the boundary of a growing monolayer of cells. As we did with the closing void, to determine if a cell is on the boundary of a tissue, we use a discrete pixel background mesh. Unlike in the previous example, a coarse mesh may be used here. We again assign both a cell-pixel interaction radius, *r*_cell-pixel_ *=* 0.55 CD, and an extended cell-pixel interaction radius, 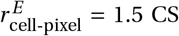, to each cell, with 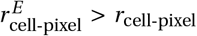. As before, if a pixel is within a cell’s cell-pixel radius, we remove the pixel. However, if a cell contains a pixel in its extended cell-pixel radius, it is labelled a boundary cell, and is part of the boundary of the tissue.

For a Voronoi tessellation with ghost nodes, a cell is labelled a boundary cell if it is neighbours with a ghost node. Lastly, for a vertex model, a cell is labelled a boundary cell if it contains a boundary labelled node. We can then check which cells are interacting with each-other to find the boundary of the tissue, defined by connected line, passing through the centre of mass of each cell. Depictions of the calculations for the OS and VT models are shown for the Fig. D7.

**Fig. D7.**
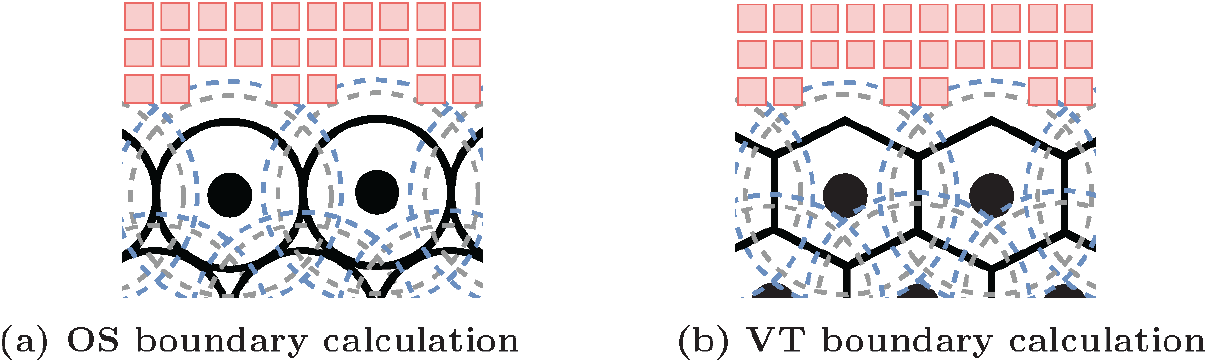
A depiction of the boundary calculation for overlapping spheres and Voronoi tessellation. The solid black dot represents the cell’s centre, the solid black circle/lines the cell’s boundary, the dashed grey circle the cell-pixel interaction radius, *r*_cell-pixel_, and the dashed blue circle the extended cell-pixel interaction radius, 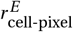. The red squares are the discrete pixel mesh

